# Independent and distinct patterns of abnormal lateral orbitofrontal cortex activity during compulsive grooming and reversal learning normalize after fluoxetine

**DOI:** 10.1101/2021.03.02.433664

**Authors:** Elizabeth E Manning, Matthew A Geramita, Sean C Piantadosi, Jamie L Pierson, Susanne E Ahmari

**Author notes:** Corresponding Author: Susanne E Ahmari. These authors contributed equally (Elizabeth E Manning, Matthew A Geramita, Sean C Piantadosi).

## Abstract

**Background:** Patients with obsessive-compulsive disorder (OCD) display disrupted performance and abnormal lateral orbitofrontal cortex (LOFC) activity during reversal learning tasks, yet it is unknown whether compulsions and reversal learning deficits share a common neural substrate. To answer this question, we measured neural activity with *in vivo* calcium imaging in LOFC during compulsive grooming and reversal learning before and after fluoxetine treatment.

**Methods:** *Sapap3*-knockout (KO) mice were used as a model for OCD-relevant behaviors. *Sapap3*-KOs and control littermates were injected with virus encoding GCaMP6f and implanted with gradient-index lenses to visualize LOFC activity using miniature microscopes. Grooming, reversal learning, and neural activity were measured pre- and post-fluoxetine treatment (18mg/kg, 4 weeks).

**Results:** Baseline compulsive grooming and reversal learning impairments in KOs improved after fluoxetine treatment. Additionally, KOs display distinct patterns of abnormal LOFC activity during grooming and reversal learning, both of which normalize after fluoxetine. Finally, modulation in response to reversal learning and compulsive behavior are independent, as reversal learning-associated neurons are distributed randomly amongst grooming-associated neurons (i.e. overlap is what would be expected by chance).

**Conclusions:** In OCD, the LOFC is disrupted during both compulsive behaviors and reversal learning, yet whether these behaviors share common neural underpinnings is unknown. We find that the LOFC plays distinct and independent roles in compulsive grooming and impaired reversal learning and their improvement with fluoxetine. These findings suggest that LOFC plays separate roles in pathophysiology and treatment of different perseverative behaviors in OCD.

## Introduction

Determining how disrupted neural activity gives rise to compulsive behaviors is vital for understanding obsessive compulsive disorder (OCD). It is commonly thought that compulsive behaviors and cognitive rigidity share a pathologic neural substrate due to an association between compulsive behavior and abnormal performance (1, 2) and reaction times (3–5) during reversal learning tasks in OCD [though see (6–8)]. Additionally, patients with OCD show abnormal activity during reversal learning (2,3,9,10) in the same orbitofrontal-striatal circuits that show abnormal activity during symptom provocation (11–13). However, it is currently unknown whether impaired reversal learning and compulsive behaviors stem from the same underlying abnormalities in orbitofrontal cortex (OFC) activity.

Determining whether the same OFC neurons show disrupted activity during both behaviors is one strategy to establish whether abnormal reversal learning and compulsive behaviors share common neural underpinnings. Since human neuroimaging technology lacks the single cell resolution necessary to test this idea, preclinical models are needed to define the precise pattern and overlap of neural activity between these distinct phenotypes. Features of the *Sapap3* knockout (KO) mouse model for OCD-relevant behaviors make it an ideal system to test this hypothesis (14–16). *Sapap3*, a post-synaptic density protein, is critical for cortico-striatal communication (14,17,18), and the *SAPAP3* gene family has been linked to OCD in candidate gene studies and in secondary analyses of genome wide association studies despite not meeting genome-wide level of significance (19–22)*. SAPAP3* transcript levels in the OFC, caudate, and nucleus accumbens are lower in patients with OCD than unaffected comparison subjects (23). Additionally, *Sapap3*- KO mice display compulsive grooming which decreases following optogenetic activation of the lateral OFC [LOFC; (15)] or treatment with a first-line pharmacotherapy for OCD, the selective serotonin reuptake inhibitor (SSRI) fluoxetine (14–16). Finally, *Sapap3*-KO mice display impaired reversal learning (24–27), strengthening their relevance to OCD.

Here, using *in vivo* microendoscopic calcium imaging in freely-moving mice, we record activity of individual LOFC neurons in *Sapap3*-KOs and wild-type (WT) littermates during grooming and reversal learning before and after 4-weeks of fluoxetine treatment. Baseline increases in compulsive grooming and impairments in reversal learning in KOs improve after fluoxetine treatment. Additionally, *Sapap3*-KOs display distinct (i.e. separate) patterns of abnormal LOFC activity during grooming and reversal learning, both of which are restored to WT levels (i.e. normalized) after fluoxetine. Finally, we show that modulation of individual LOFC neurons in response to reversal learning and compulsive behavior is independent, as the overlap between reversal learning-associated neurons and grooming-associated neurons is what would be expected by chance. Consequently, disrupted LOFC modulation in response to reversal learning is distinct from and independent of disrupted modulation in response to compulsive grooming in *Sapap3*-KO mice.

## Methods

### Animals

All procedures were carried out in accordance with the guidelines for care and use of laboratory animals from the NIH and with approval from the University of Pittsburgh Institutional Animal Care and Use Committee (IACUC). *Sapap3*-KOs (n = 8; 5 female) and WT (n = 6; 3 female) littermates were maintained on a C57BL/6 background. Mice were group-housed with 2–5 same-sex mice per cage in reverse light cycle (12:12, lights on at 7pm) with *ad libitum* access to food and water, except during operant training when food was restricted. All tests were conducted under red or infrared lighting during dark cycle.

### Calcium imaging surgery

Mice underwent two surgeries for optical imaging studies similar to standard protocols (28, 29). For the first surgery, 800nl of a virus encoding GCaMP6f under the synapsin promotor (AAV5-synapsin-GCaMP6f-WPRE-SV40, titer 1.82×10^12^; Penn Vector Core) was injected into the LOFC (AP: +2.7, ML: -1.0, DV: -1.6), followed by implantation of a gradient refractive index lens (ProView GRIN lens, 6mm long x 0.5mm wide; Inscopix, Palo Alto, CA USA) dorsal to the viral injection target (AP: +2.6, ML: -1.2, DV: -1.4). After 3-4 weeks of virus expression, a magnetic microscope baseplate (Inscopix) was implanted to allow attachment of the miniature microscope (nVista 2.0, Inscopix). More details can be found in the Supplement. Lens placements for individual animals are shown in Fig.S1. Based on analysis of calcium event rates and widths, over 90% of imaged neurons are putative excitatory neurons (Fig.S2; see Supplemental Discussion).

### Fluoxetine administration

(±)Fluoxetine hydrochloride (Fluoxetine; NIMH Chemical Synthesis and Drug Supply Program) was administered via drinking water according to established methods (30). 100mg/L fluoxetine hydrochloride was administered for 4 weeks to achieve a target dose of 18mg/kg, followed by a 2- week drug washout period to help determine whether changes in behavior are due to fluoxetine or additional training (31).

### Experimental design

Behavior and neural data were collected weekly for grooming and every other week for reversal learning (Fig.1b). In weeks where both behaviors were tested, grooming assessment was conducted first, and mice underwent operant training for 2 days prior to reversal imaging on the third day (details below). Neural and behavior data are presented from baseline and 4-week treatment sessions, and behavior data is also presented from the reversal learning washout session.

**Figure 1:**
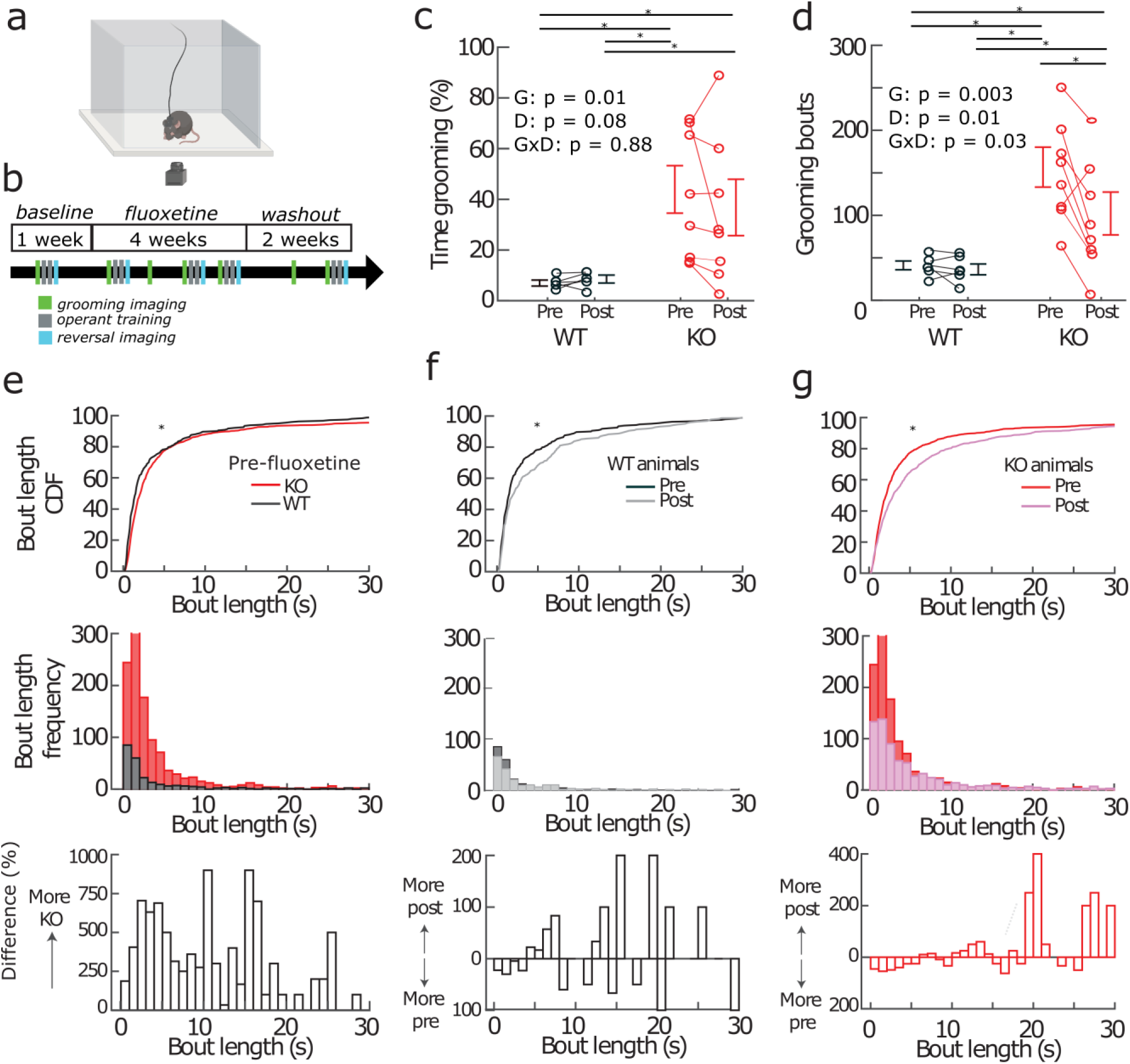
Baseline increases in the number of grooming bouts, but not total time grooming, improve after fluoxetine treatment in KOs. (a) Grooming was tested in a clear acrylic chamber positioned above a behavior acquisition camera. (b) Experimental timeline: Baseline grooming (green) and reversal learning (cyan) were measured prior to 4 weeks of fluoxetine treatment. While fluoxetine was administered, grooming was assessed weekly and reversal learning was assessed at weeks 1,3,4. Post-fluoxetine data presented throughout the manuscript are from the 4-week time point. Grooming and reversal learning were also assessed after a 2 week washout period. (c) Data collected from 8 KO mice (red) and 6 WT littermates (black). Both pre- and post-fluoxetine, KOs spent more time grooming than WTs (genotype: F_(1,12)_ = 8.5,p = 0.01; drug: F_(1,12)_ = 3.6, p = 0.08; genotype x drug: F_(1,12)_ =0.02, p = 0.88). (d) Pre-fluoxetine increases in the number of grooming bouts in KOs decrease after fluoxetine (genotype: F_(1,12)_ =13.5, p = 0.003; drug: F_(1,12)_ = 8.8, p = 0.01; genotype x drug: F_(1,12)_ = 6.0, p = 0.03). (e-g) Differences in bout length between (e) WTs and KOs prior to fluoxetine treatment; (f) WTs pre- and post-fluoxetine; and (g) KOs pre- and post-fluoxetine. *Top*: Cumulative distribution function (CDF) of bout length. *Middle*: Histogram of bout length. *Bottom*: Percent difference in bout length frequency. (e) Bout lengths are longer in KOs compared with WTs (KS – p < 0.05). (f-g) Fluoxetine increases the length of grooming bouts in WTs (e; KS – p < 0.05) and KOs (f; KS – p < 0.05). P-values of the repeated measures ANOVA are indicated in the inset statistics such that G: genotype, D: drug, GxD: interaction. Asterisks in panels c-d indicate post-hoc t-tests with p-values that are less than 0.05.

### Grooming imaging procedures

Mice were tested in a clear acrylic chamber (8”x8”x12”) positioned above a behavior acquisition camera (Fig.1a; Point Grey Blackfly, FLIR Integrated Imaging Solutions, 40Hz). Behavior video and calcium signals were synchronized by a central data acquisition box (LabJack U3-LV, Labjack Corporation, Lakewood CO USA). Neural and behavior data were recorded for 40 minutes. Videos were analyzed offline by a trained experimenter blind to genotype and treatment using Observer XT software (Noldus, Leesburg, VA). Frame-by-frame analysis was used to identify the start and end of face and body grooming and hind-leg scratching (more details available in Supplement).

### Operant conditioning imaging procedures and analysis

Mice were tested in a reversal learning paradigm using operant chambers (Med Associates, Fairfax, VT) similar to previously described (*24*) with modifications for imaging studies (see Supplement). Mice were pretrained on the task, and for each timepoint they were tested for three days. The first two days used the same rule (e.g. left lever correct, right lever incorrect, using the contingency from their most recent prior test), and on the third day the contingency was reversed (e.g. right lever correct) and calcium imaging was performed. Mice were trained on a variable ratio (VR) 2 schedule, and rewarded correct responses resulted in retraction of the two levers and delivery of a reward pellet. Operant behavior events and calcium signals were synchronized via TTL pulses from MED-PC system and frame information from Inscopix sent directly to a central data acquisition box (Labjack). Neural data and behavior were recorded for 30 minutes but only the first 15 minutes were analyzed (see Supplement). Time to acquire the correct lever press was estimated by the mid-point of the sigmoid (1) fit to the cumulative distribution of correct lever presses during the first 15 minutes. In the formula, A and B are the slope and midpoint of the sigmoid, respectively.

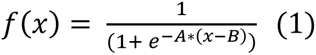

### Imaging acquisition, processing and analysis

nVistaHD software recorded fluorescent signal as compressed greyscale tiffs (20Hz, 470nm LED power 0.1-0.3mW, image sensor gain = 1-4). All imaging pre-processing was performed using Mosaic software (version 1.2.0, Inscopix) via custom Matlab (MATHWORKS, Natick MA, USA) scripts. Data were decompressed, downsampled (x4 spatial and x2 temporal), and motion corrected before fluorescent signals were extracted from individual neurons using Constrained Nonnegative Matrix Factorization for Endoscopic data (CNMFe) (32), with putative neurons manually sorted by blind observer (28).

### Longitudinal tracking of neurons across sessions

Putative neurons identified via CNMFe were matched across sessions using *CellReg* (33). More details are available in the Supplement.

### Encoding model

To determine the extent to which LOFC neurons were modulated in response to grooming or reversal learning events, we modified a previously described multiple linear regression encoding model (34). For details see Supplement.

### Statistical analysis

Effects of genotype and fluoxetine on behavior were assessed using repeated measures ANOVA. Percentages of cells modulated and strength of modulation were assessed using repeated measures ANOVA and linear regression, respectively. Where interactions were observed in ANOVAs, post hoc t tests were performed (details and p values in table 3). Reversal learning imaging metrics were adjusted for differences in the number of behavioral events (see Supplement). Description of analysis of expected versus actual overlap between grooming and reversal learning can be found in the Supplement. Graphs show individual datapoints and SEM unless otherwise stated; mean and SEM are available in table 3.

## Results

### Analysis of grooming behavior

*Sapap3*-KOs groomed for a larger percentage of time both at baseline and after 4 weeks of fluoxetine treatment (Fig.1c; genotype: F_(1,12)_ = 8.5,p = 0.01; drug: F_(1,12)_ = 3.6, p = 0.08; genotype x drug: F_(1,12)_ =0.02, p = 0.88). In contrast, baseline increases in the number of grooming bouts in KOs decreased after fluoxetine treatment (Fig.1d; genotype: F_(1,12)_ =13.5, p = 0.003; drug: F_(1,12)_ = 8.8, p = 0.01; genotype x drug: F_(1,12)_ = 6.0, p = 0.03). KOs engaged in longer grooming bouts compared to WTs, which was apparent across all bout lengths (Fig.1e; p < 0.001). The number of bouts, but not the total time grooming, decreased after fluoxetine because the length of grooming bouts significantly increased after fluoxetine in both WTs (Fig.1f) and KOs (Fig.1g).

### Neural activity during grooming

To determine whether LOFC neurons were modulated during grooming, we analyzed *in vivo* calcium imaging data (Fig.2a-d) by adapting a previously described linear regression model (34) (see Supplement; Fig.2e). This approach allowed us to quantify how LOFC neurons were modulated in response to grooming using two different metrics: 1) the percentage of LOFC neurons that were significantly modulated in response to grooming, and 2) the strength of modulation in each significant neuron. First, we assessed the total population of cells (Fig.2f; Table 1). In KOs, baseline increases in the percentage of LOFC neurons inhibited during grooming (groom_inh_) normalized after fluoxetine (Fig.2g; genotype: F_(1,12)_ = 6.90, p = 0.02; drug: F_(1,12)_ > 100, p < 0.001; drug x genotype: F_(1,12)_ > 100, p < 0.001). Both at baseline and after fluoxetine, similar percentages of LOFC neurons from KOs and WTs were excited during grooming (groom_exc_) (Fig.2h). Additionally, both groom_inh_ (Fig.2i) and groom_exc_ (Fig.2j) neurons from KOs were modulated in response to grooming more strongly than neurons from WTs both at baseline and after fluoxetine. Interestingly, while the percentage of groom_inh_ neurons did not correlate with the number of grooming bouts at baseline in KOs or WTs (Fig.2k), the fluoxetine-associated decrease in the percentage of groom_inh_ neurons significantly correlated with a decrease in the number of grooming bouts in KOs but not WTs (Fig.2l; KO: r = 0.80, p = 0.01; WT: r = 0.40, p = 0.41). Consequently, these analyses suggest that decreases in grooming in KOs after fluoxetine are due, in part, to decreases in the percentage of groom_inh_ LOFC neurons.

**Figure 2:**
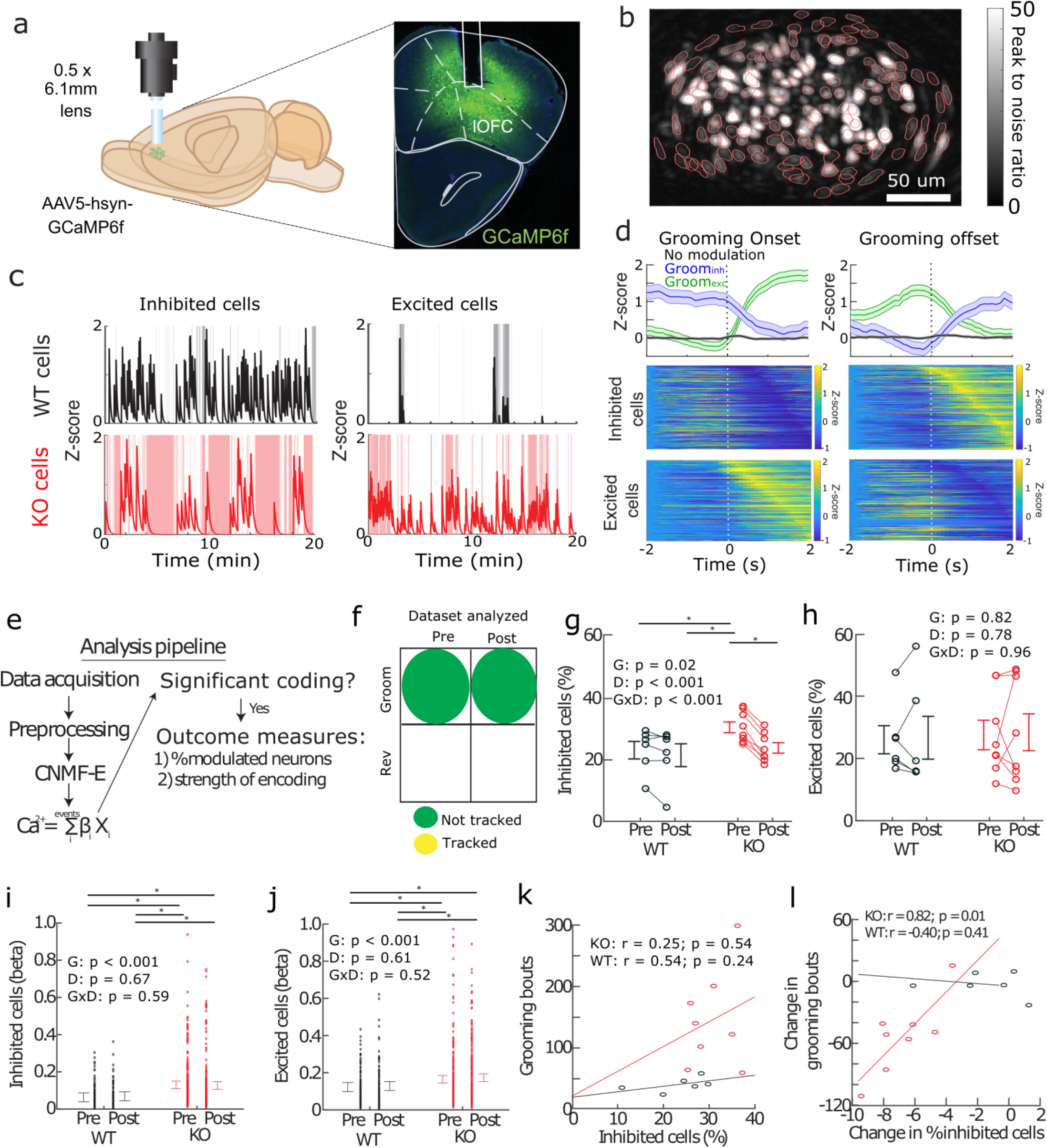
Fluoxetine-associated decreases in the number of grooming bouts are correlated with a decrease in the percentage of cells inhibited by grooming in KOs. (a) *Left:* Schematic of calcium imaging in the LOFC. *Right:* Representative coronal section showing lens placement and GCaMP6f expression. (b) Individual contours of putative LOFC neurons (red) plotted on top of the peak-to-noise ratio image of cell activity (scale bar=50um). (c) Examples of cells that are most strongly modulated in response to grooming from each genotype (grooming periods are depicted by grey/red shaded background). (d) (*Top*): Calcium traces averaged across all groom_exc_ (green), groom_inh_ (blue), and not modulated (black) neurons in all animals aligned to the onset (*Left*) or offset (*Right*) of grooming bouts. (*Bottom*): Heatmaps for all inhibited (*Top*) or excited (*Bottom*) neurons modulated in response to grooming aligned to onset (*Left)* or offset (*Right*). (e) Analysis pipeline: After data were acquired and preprocessed, calcium transients from individual cells were identified using CMNFe. Presence of significant modulation in response to behavior was determined using encoding model. Two measures of modulation were used: 1) the percent of neurons that were significantly modulated in response to the behavior and 2) the strength of modulation of individual neurons that are significantly modulated (See Methods). (f): Cells are not tracked between pre- and post-fluoxetine sessions in this analysis. (g-h) Percentage of cells modulated in response to grooming pre- and post-fluoxetine in WTs and KOs. (g) Pre-fluoxetine increases in the percentage of inhibited neurons in KOs decreased after fluoxetine (genotype: F_(1,12)_ = 6.90, p = 0.02; drug: F_(1,12)_ > 100, p < 0.001; drug x genotype: F_(1,12)_ > 100, p < 0.001). (h) There were no genotype or fluoxetine effects on the percentage of excited cells (genotype: F_(1,12)_ = 0.05, p = 0.82; drug: F_(1,12)_ = 0.08, p = 0.78; genotype x drug: F_(1,12)_ = 0.003, p = 0.96). (i-j) Beta weights for cells significantly modulated in response to grooming pre- and post-fluoxetine in WTs and KOs. Increased modulation strength by grooming in KOs does not change after fluoxetine in (i) inhibited cells (genotype: p < 0.001; drug: p = 0.67; genotype x drug: p = 0.59) or (j) excited cells (genotype: p < 0.001; drug: 0.61; genotype x drug: p = 0.52). (k) No correlation between baseline number of grooming bouts and the percentage of inhibited cells. (l) Decreases in the number of grooming bouts after fluoxetine were correlated with decreases in the percentage of inhibited cells in KOs, but not WTs. P-values of the repeated measures ANOVA are indicated in the inset statistics such that G: genotype, D: drug, GxD: interaction. Asterisks in panels g-j indicate post-hoc t-tests with p-values that are less than 0.05.

**Table 1:** Number of cells across mice.

| Animal | Genotype | Pre Grooming | Post Grooming | Pre Reversal | Post Reversal |
| --- | --- | --- | --- | --- | --- |
| 3001 | WT | 128 | 117 | 131 | 109 |
| 3297 | WT | 57 | 68 | 77 | 89 |
| 3298 | WT | 111 | 84 | 108 | 91 |
| 3592 | WT | 106 | 72 | 70 | 101 |
| 3604 | WT | 137 | 103 | 133 | 107 |
| 3632 | WT | 109 | 112 | 62 | 118 |
| Avg ± SEM | WT | 108 ± 28 | 93 ± 21 | 97 ± 31 | 103 ± 11 |
| 2991 | KO | 127 | 134 | 122 | 123 |
| 2993 | KO | 116 | 131 | 71 | 109 |
| 3091 | KO | 180 | 107 | 84 | 141 |
| 3605 | KO | 124 | 129 | 106 | 149 |
| 3616 | KO | 153 | 136 | 108 | 142 |
| 3617 | KO | 133 | 139 | 144 | 123 |
| 3626 | KO | 88 | 64 | 74 | 69 |
| 3627 | KO | 81 | 38 | 59 | 56 |
| Avg ± SEM | KO | 125 ± 32 | 110 ± 38 | 96 ± 29 | 114 ± 34 |

Decreases in the percentage of groom_inh_ LOFC neurons after fluoxetine treatment could arise in two ways – fewer neurons could remain inhibited and/or fewer new neurons could become inhibited. To test these two possibilities, the same neurons were tracked across grooming sessions (Fig.3a; see Methods; Table 2). Much like the untracked dataset (Fig.2), there was a significantly higher baseline percentage of groom_inh_ neurons in KOs that decreased after fluoxetine treatment (Fig.3b,c). While there were no genotype differences in the percentage of neurons that *remained* inhibited during grooming after fluoxetine (Fig.3d), fewer LOFC neurons *became* newly inhibited during grooming in KOs (Fig.3e; p < 0.001). Therefore, fluoxetine-associated decreases in the percentage of groom_inh_ neurons in KOs are partly due to fewer neurons becoming inhibited after fluoxetine treatment.

**Figure 3:**
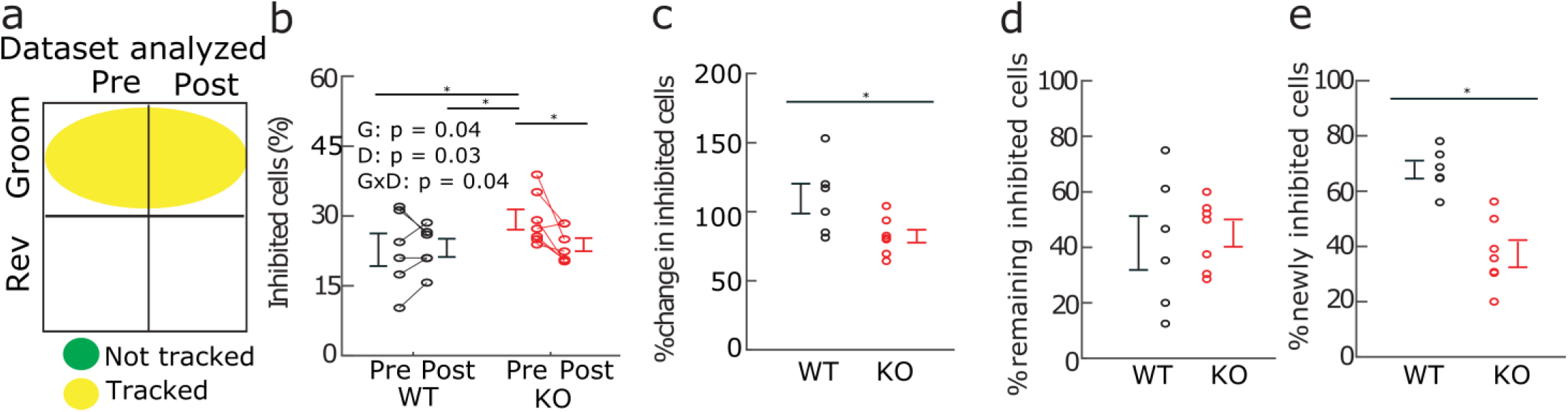
Decreased percentage of inhibited cells in KOs is due to fewer newly inhibited cells rather than fewer cells that remain inhibited after fluoxetine. (a) Cells tracked between pre- and post-fluoxetine grooming sessions. (b) Similar to the untracked dataset (Fig. 2), KOs show an increased percentage of grooming inhibited neurons that decreases after fluoxetine (genotype: F_(1,12)_ = 5.03, p = 0.04; drug: F_(1,12)_ = 5.9, p = 0.03; genotype x drug: F_(1,12)_ = 5.1, p = 0.04). (c) The percent change in inhibited cells is lower in KOs (t-test: p = 0.02). Percent change in inhibited cells is calculated as the difference in inhibited cells between pre- and post-fluoxetine divided by the inhibited cells pre-fluoxetine. (d) The percentage of neurons that remain inhibited by grooming after fluoxetine does not differ by genotype (t-test: p = 0.54). (e) Fewer new neurons become inhibited by grooming after fluoxetine in KOs compared with WTs (t-test: p < 0.001). Asterisks in panel (b) indicate post-hoc t-tests with p-values that are less than 0.05.

**Table 2:**
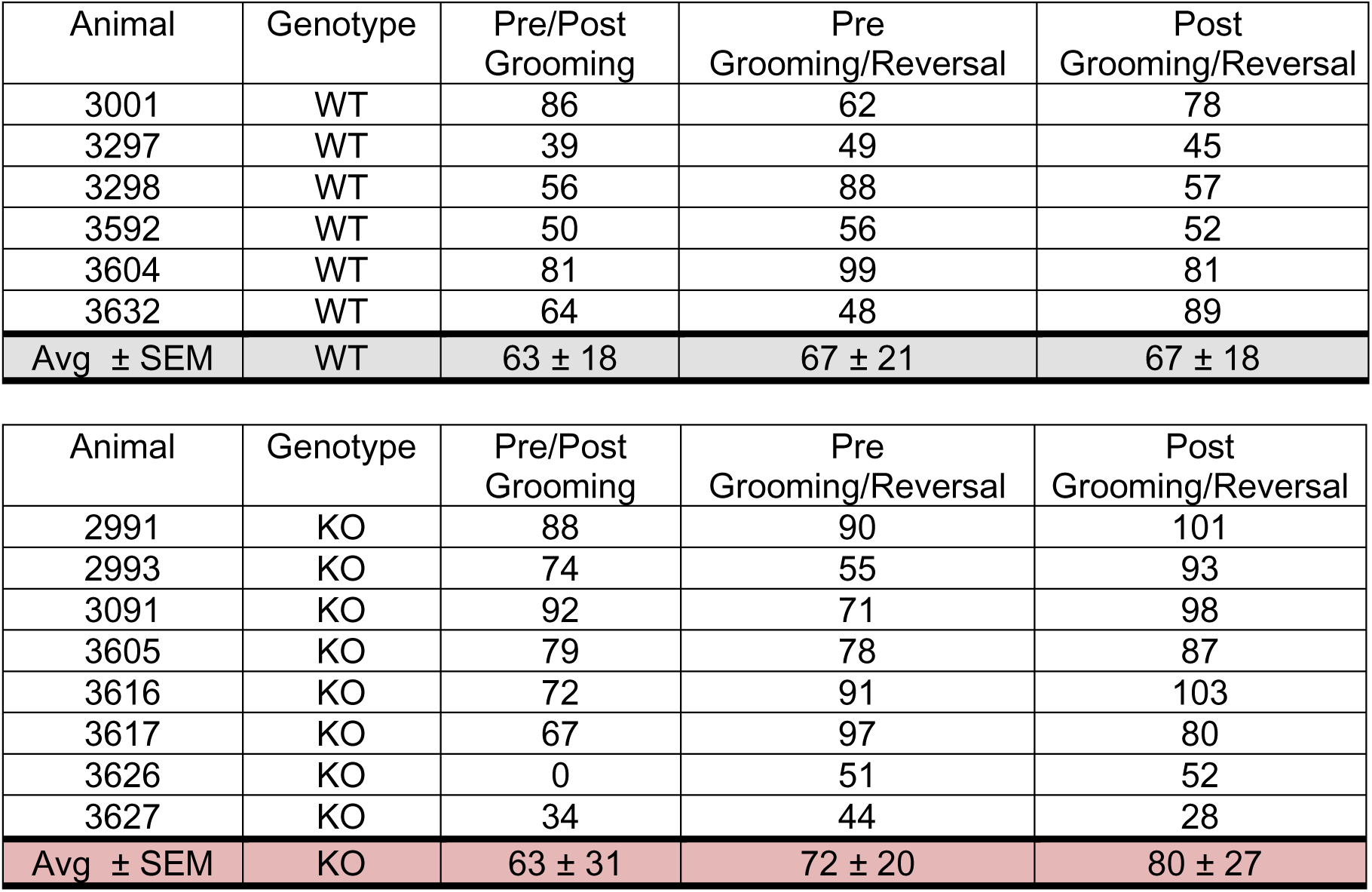
Number of tracked cells across mice.

### Analysis of reversal-learning behavior

We next assessed the patterns of neural activity associated with reversal learning. Mice were first trained to associate one of two levers with reward on a VR2 schedule as previously described (24) (see Methods; Figs.4a,b). No genotype differences were observed in pre-reversal discrimination (Fig.S3). Fluoxetine normalized baseline reductions in the number of correct lever presses (Fig.4c; genotype: F_(1,12)_ = 5.72, p = 0.03; drug: F_(1,12)_ = 0.76, p = 0.39; genotype x drug: F_(1,12)_ = 4.55, p = 0.04) and increases in time to acquire the correct lever (see Methods; Fig.4d,e; genotype: F_(1,12)_ = 10.01, p = 0.008; drug: F_(1,12)_ = 1.31,p = 0.25; genotype x drug: F_(1,12)_ = 6.87, p = 0.02) in KOs. Since correct lever presses trigger reward delivery and the start of the next trial, significant baseline reductions in the number of rewards received (Fig.4f) and trials completed (Fig.S4a-c) also normalized after fluoxetine in KOs. Additionally, KO and WT mice showed similar significant decreases in incorrect lever presses after fluoxetine (Fig.4g). No genotype- or fluoxetine- associated differences in the number of total (Fig.S4d-f) or unrewarded (Fig.S4g-i) magazine entries were observed.

**Figure 4:**
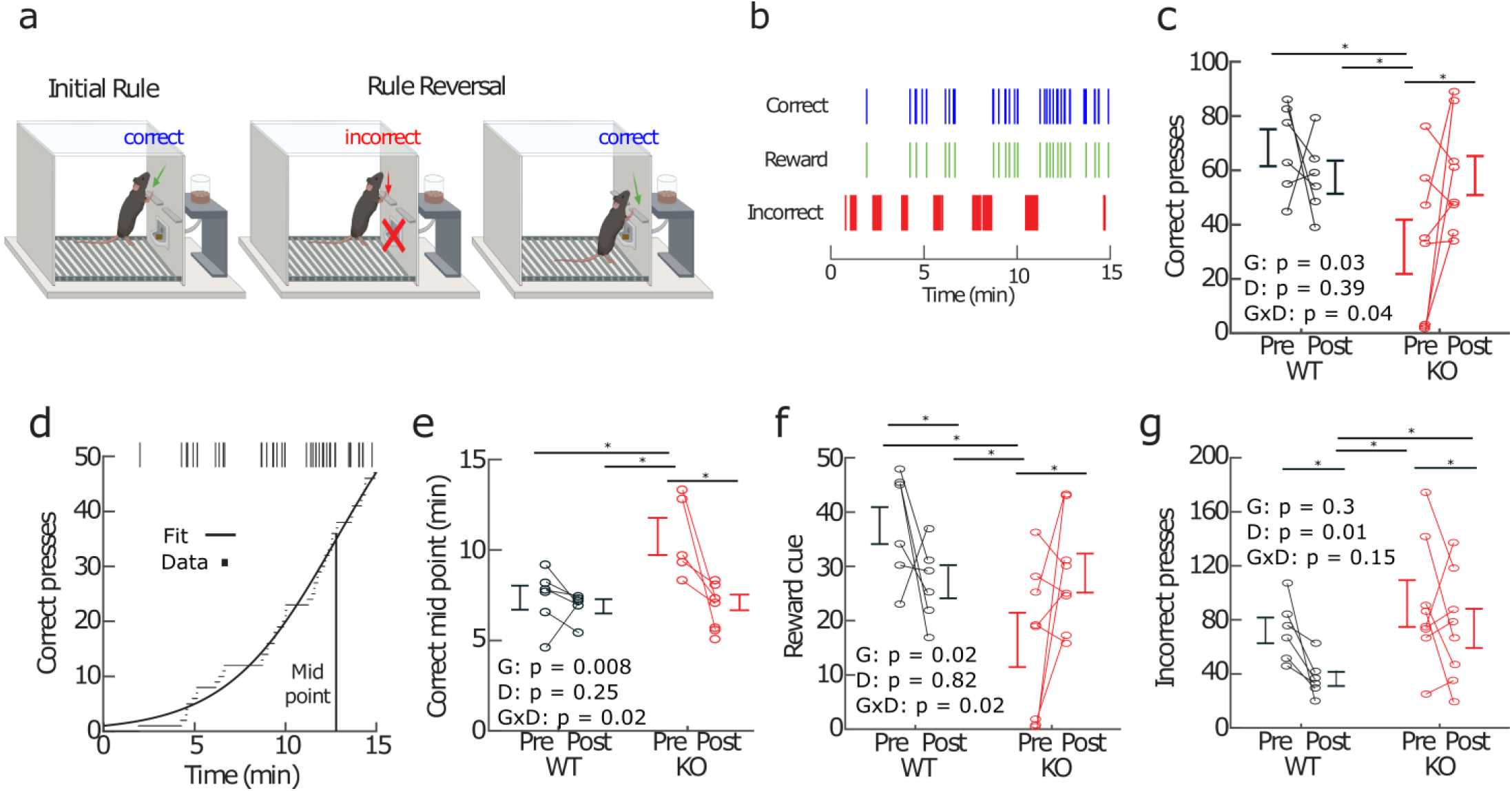
Baseline deficits in reversal learning in KOs improve after fluoxetine treatment. (a) Schematic of reversal learning task design. During operant training, mice associate one lever with reward. After reversal of lever contingencies, mice must learn that the other lever now delivers reward. (b) Example event raster of relevant events in the reversal learning task for one animal. (c) Decreases in the number of correct lever presses in KOs normalize after fluoxetine (genotype: F_(1,12)_ = 5.72, p = 0.03; drug: F_(1,12)_ = 0.76, p = 0.39; genotype x drug: F_(1,12)_ = 4.55, p = 0.04). (d) Example of fitting sigmoid to CDF of correct lever presses. The mid-point of the sigmoid is used as a measure of the time to acquire correct lever presses (See Methods). The 3 KOs that did not learn to press the correct lever (panel c) are excluded from the pre-fluoxetine mid-point calculation. (e) Increases in the time to acquire the correct lever decrease after fluoxetine in KOs (genotype: F_(1,12)_ = 10.01, p = 0.008; drug: F_(1,12)_ = 1.31,p = 0.25; genotype x drug: F_(1,12)_ = 6.87, p = 0.02). (f) Because correct lever presses trigger rewards, decreases in the number of rewards in KOs also increase after fluoxetine (genotype: F_(1,12)_ = 6.88, p = 0.02; drug: F_(1,12)_ = 0.04, p = 0.82; genotype x drug: F_(1,12)_ = 6.59, p = 0.02). (g) The number of incorrect lever presses decreases after fluoxetine in both genotypes (genotypes: F_(1,12)_ = 1.19, p = 0.30; drug: F_(1,12)_ = 8.87, p = 0.01; genotype x drug: F_(1,12)_ = 2.31, p = 0.15). P-values of repeated measures ANOVA are indicated in the inset statistics such that G: genotype, D: drug, GxD: interaction. Asterisks in panels (c-g) indicate post-hoc t-tests with p-values that are less than 0.05.

To determine whether differences in behavior following 4 weeks of fluoxetine treatment were due to drug versus additional training on the task, reversal learning was repeated after a 2-week fluoxetine washout period. After washout, the number of correct lever presses significantly decreased (Fig.S5a-c) and the time to acquire correct presses significantly increased (Fig.S5d-f), indicating prior improvements in reversal learning were due to fluoxetine. In contrast, decreases in the number of incorrect lever presses were still observed after washout in both genotypes (Fig.S5j-l), suggesting that these changes were likely due to additional training. Together, these analyses indicate that baseline behavioral deficits in reversal learning in KOs normalized due to fluoxetine treatment.

### Neural activity during reversal learning

We next determined how LOFC neurons were modulated in response to the reversal learning task using a variant of the linear regression model described above. In this model, five behavioral variables – three actions (correct lever presses, incorrect lever presses and magazine entries) and two sensory stimuli (reward cues and trial start cues) – were used to predict each cell’s calcium signal (Figs.5a-d). Notably, there were no genotype- or fluoxetine-associated differences in the percentage of cells excited in response to any of the five behavioral variables (Fig.5e-g, Fig.S6a,b) (few neurons were inhibited by task events and consequently were not included in these analyses; Fig.S7). In contrast, baseline deficits were observed in the strength of modulation in response to both correct lever presses (Fig.5h; genotype: p = 0.01; drug: p = 0.18; genotype x drug: p = 0.04) and the reward cue (Fig.5i; genotype: p = 0.001; drug: p = 0.45; genotype x drug: p = 0.01) in KOs; these deficits were normalized after fluoxetine treatment. Modulation in response to correct lever presses and reward cue and changes following fluoxetine did not correlate with the time to acquire correct lever presses after reversal (Fig.S8). Additionally, deficits in modulation in response to the reward cue were unlikely to be due to a generalized sensory processing deficit as there was no genotype- or fluoxetine-associated difference in modulation in response to the trial start cue (Fig.S6c). The strength of modulation in response to the incorrect lever press (Fig.5j) significantly decreased after fluoxetine treatment in both WTs and KOs. Finally, there were no genotype- or fluoxetine-associated differences in the modulation in response to magazine entries (Figs.S6d). Therefore, fluoxetine-associated improvements in reversal learning in KOs were associated with improvements in the strength of modulation in response to correct lever presses and the reward cue.

**Figure 5:**
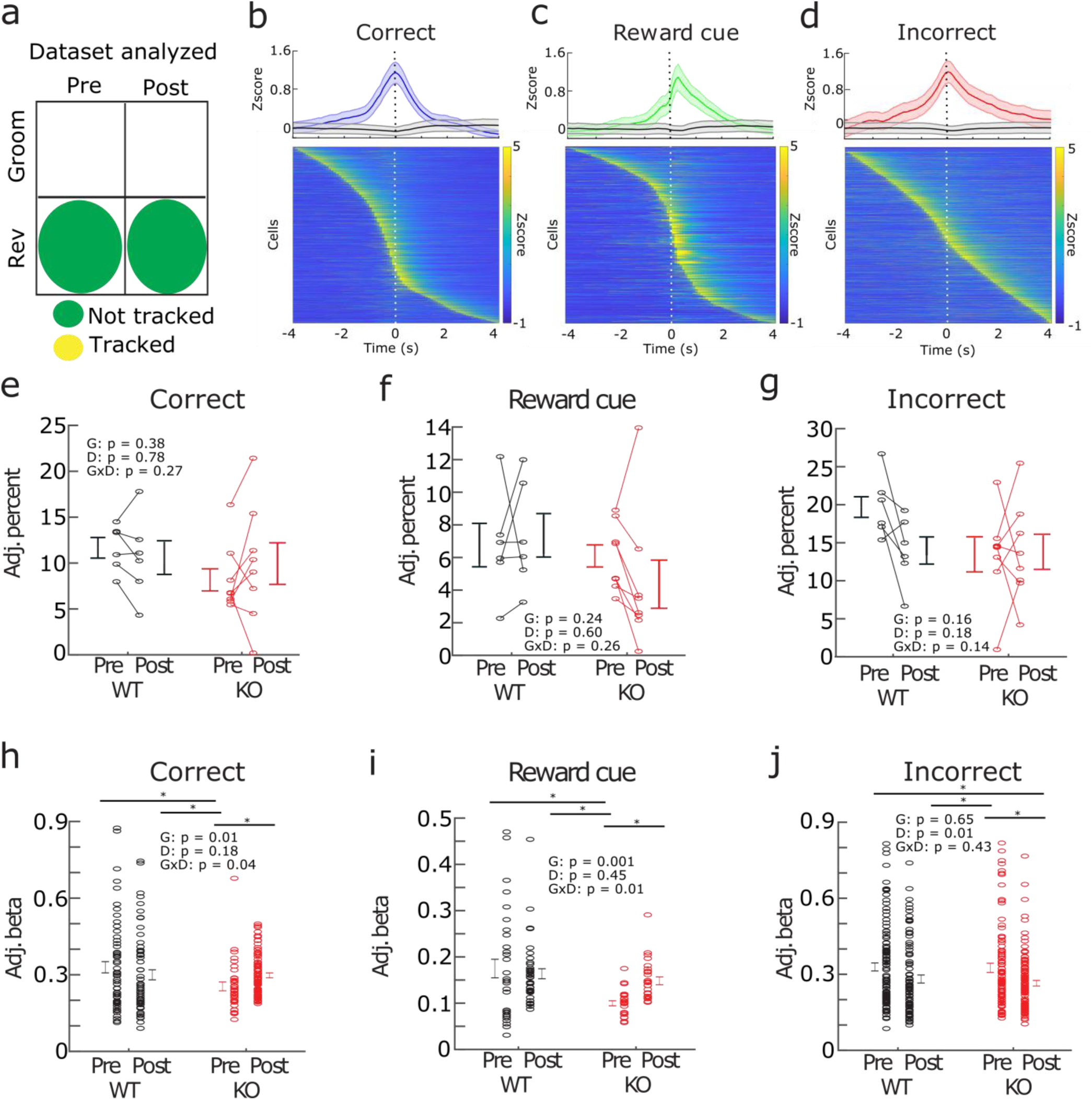
Baseline deficits in the strength of modulation in response to correct lever press and reward cue in LOFC neurons from KOs normalize after fluoxetine treatment. (a) Cells taken from the untracked dataset. (b) *Top:* Average z-scored calcium trace from all neurons combined from all animals that are significantly modulated in response to correct lever press (blue) and not significantly modulated in response to the correct lever press (grey). Calcium transients are time locked to correct lever presses (time 0). *Bottom:* Heatmap of calcium activity time-locked to correct lever presses from all cells significantly modulated in response to correct lever presses across all animals. Cells are sorted according to max activity at time 0. (c) Same as (b) but for cells significantly modulated in response to reward cues (green). (d) Same as (b) but for cells significantly modulated in response to incorrect lever presses (red). (e-g) Percentages of modulated neurons were adjusted for the differences in the number of task events (“Adj. percent”). Plots display residuals after correcting for differences in the number of behavioral events. There were no genotype or fluoxetine effects on the percentage of neurons modulated in response to (e) correct lever presses (genotype: F_(1,12)_ = 0.77, p = 0.38; drug: F_(1,12)_ = 0.075, p = 0.78; genotype x drug: F_(1,12)_ = 1.3, p = 0.27), (f) reward cues (genotype: F_(1,12)_ = 1.49, p = 0.24; drug: F_(1,12)_ = 0.29, p = 0.60; genotype x drug: F_(1,12)_ = 1.34, p = 0.26), or (g) incorrect lever presses (genotype: F_(1,12)_ = 2.24, p = 0.16; drug: F_(1,12)_ = 2.02, p = 0.18; genotype x drug: F_(1,12)_ = 2.50 p = 0.14). (h-j) Genotype and fluoxetine effects on the strength of modulation in response to task variables in significantly modulated neurons. Variables adjusted for number of task events and animal-to-animal variability (“Adj. beta”). Fluoxetine normalizes baseline reductions in the strength of modulation in response to (h) correct lever presses (genotype: p = 0.01; drug: p = 0.18; genotype x drug: p = 0.04) and (i) reward cues (genotype: p = 0.001; drug: p = 0.45; genotype x drug: p = 0.01). (j) The strength of modulation in response to incorrect lever presses decreases after fluoxetine in both genotypes (genotype: p = 0.65; drug: p = 0.01; genotype x drug: p = 0.43). P-values of repeated measures ANOVA are indicated in the inset statistics such that G: genotype, D: drug, GxD: interaction. Asterisks in panels (e-j) indicate post-hoc t-tests with p-values that are less than 0.05.

### Interaction of LOFC neural activity during grooming and reversal learning

We next sought to determine how compulsive grooming and reversal learning behaviors interact. We first examined our behavioral data. Performance on the reversal learning task did not correlate with grooming levels, similar to our prior observations (24) and those of other groups (25, 27) (Fig.S9). Next, we assessed the overlap between neurons that were modulated in response to reversal learning and neurons that were modulated in response to compulsive grooming. LOFC neurons were tracked between grooming and reversal learning sessions at both the pre- and post- fluoxetine time points and pooled across animals of the same genotype to improve statistical power (see Supplement; Figs. 6a,f; Table 2). To determine whether the actual overlap between reversal learning and grooming-modulated cells was different from that which would be expected by chance, we performed a bootstrap analysis (see Supplement). For both pre- and post-fluoxetine, the actual overlap between cells modulated in response to the five reversal learning parameters and grooming fell within the 95% confidence intervals of the overlap expected by chance for nearly all comparisons except for overlap between groom_inh_ neurons and neurons modulated in response to the incorrect press pre-fluoxetine (Figs.6c-e,h-j, Figs.S10). These data indicate that cells modulated in response to reversal learning are randomly distributed among cells modulated in response to grooming (Fig.8; e.g. are overlapping and/or segregated at the level expected by chance).

**Figure 6:**
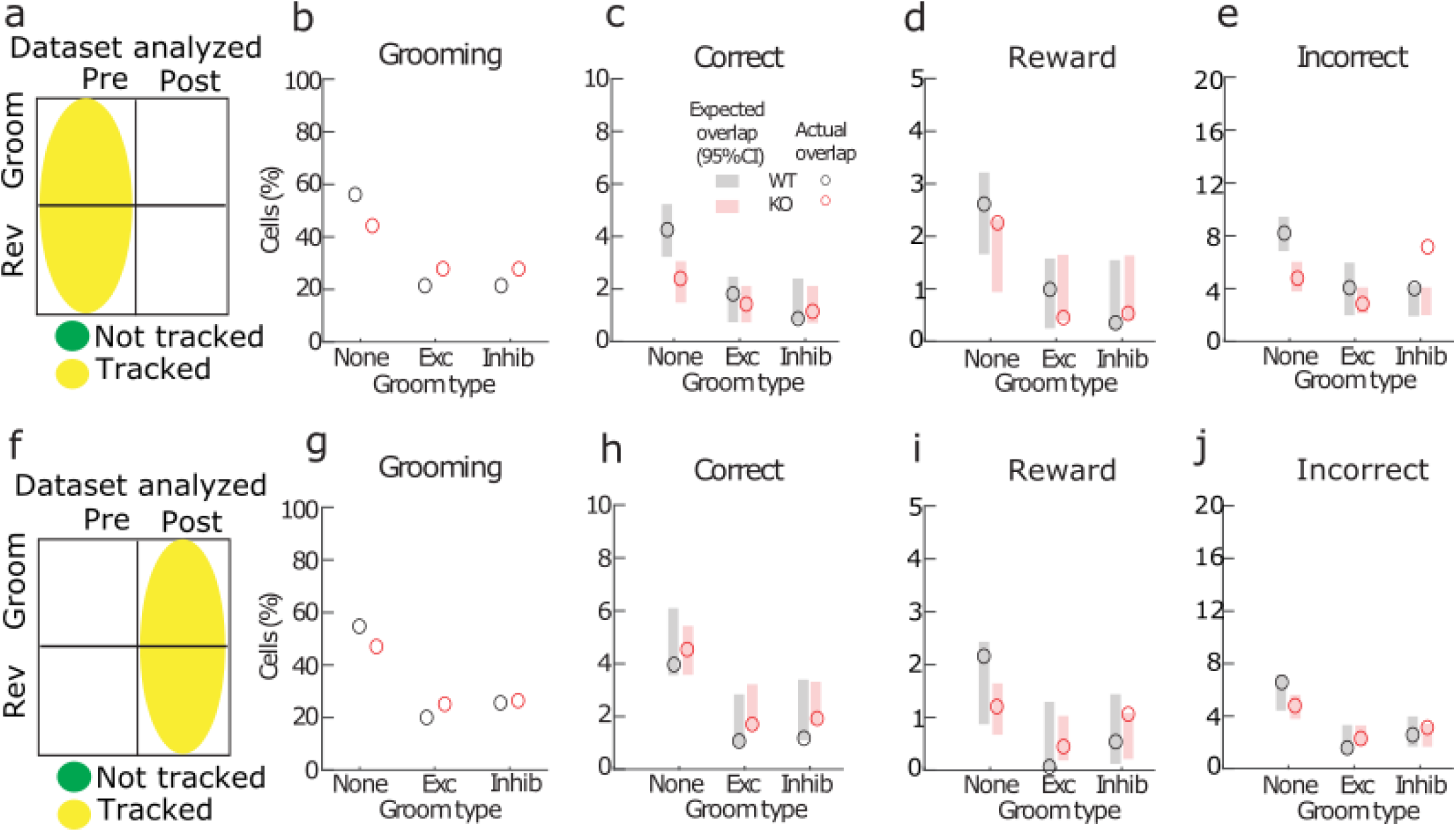
Reversal learning LOFC neurons are randomly distributed among grooming cells. (a) Cells were tracked between grooming and reversal sessions prior to fluoxetine. (b) Percentage of cells modulated in response to grooming after pooling of all grooming cells across animals of the same genotype (WT black; KO red). Modified bootstrap analysis was performed (See Methods) to determine the expected overlap between populations modulated in response to reversal learning and grooming. Populations were considered significantly overlapping or segregated if the actual overlap (circles) was above or below the 95% CI of the expected overlap (shaded rectangles), respectively. (c-e) Reversal learning cells are randomly distributed among grooming cells. Neurons modulated in response to (b) correct lever presses, (d) reward cues, and (e) incorrect lever presses are randomly distributed among grooming cells. The only exception is a greater than expected overlap between incorrect neurons and those inhibited by grooming. (f-j) Same as a-e but for cells tracked between post-fluoxetine grooming and reversal sessions.

Finally, we analyzed whether modulation in response to grooming influenced the strength of modulation in response to reversal learning. Similar to the untracked dataset (Fig.5), tracked neurons in KOs (Fig.7a-c) showed deficits in the strength of modulation in response to both correct lever presses (Fig.7d) and reward cues (Fig.7g), which normalized after fluoxetine. Pre-fluoxetine, genotype differences in the strength of modulation were not influenced by whether a neuron was modulated in response to grooming, for either correct lever presses (Fig.7e; genotype: p < 0.001; groom modulation: p = 0.01; genotype x groom mod: p = 0.56) or reward cues (Fig.7h; genotype: p = 0.1; groom modulation: p = 0.04; genotype x groom mod: p = 0.85). Surprisingly, both pre- and post-fluoxetine, groom_inh_ neurons from both WTs and KOs showed weaker modulation in response to correct lever presses (Fig.7e,f) and reward cues (Fig.7h,i) than neurons that were not modulated in response to grooming. Similarly, groom_inh_ neurons were also modulated in response to magazine entries more weakly than neurons that were not modulated in response to grooming in both genotypes (Fig.S11a-c). There were no effects of grooming-modulation on modulation in response to incorrect lever presses (Fig.S11d-f) or trial start cues (Fig.S11g-i). Therefore, degraded modulation in response to correct lever presses, reward cues and magazine entries in groom_inh_ neurons is independent of genotype and fluoxetine treatment. Thus, these results suggest that whether or not a neuron is modulated in response to grooming does not contribute to pathologic deficits in the strength of modulation in response to reversal learning in KOs.

**Figure 7:**
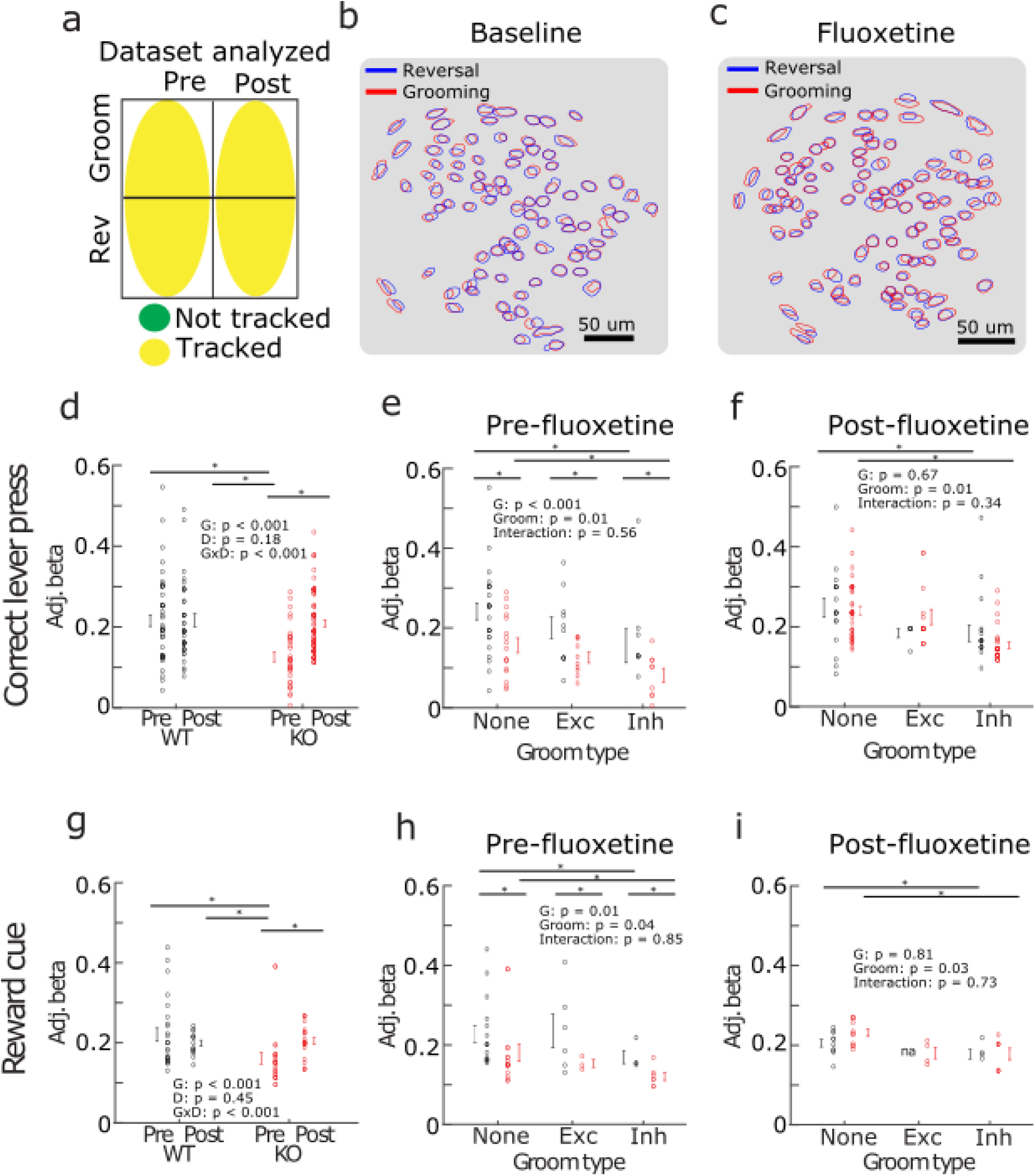
Presence of grooming modulation does not contribute to weaker modulation in response to reversal learning in KOs. (a) Cells are tracked between grooming and reversal sessions at both the pre- and post-fluoxetine time points. (b-c) Contour maps of putative LOFC neurons from a representative mouse aligned across reversal (blue) and grooming (red) sessions at baseline (left) and after 4 weeks of fluoxetine treatment (right). (d) Similar to Figure 5g, baseline deficits in modulation in response to correct lever presses normalize after fluoxetine in KOs (genotype: p < 0.001; drug: p = 0.18; genotype x drug: p < 0.001). (e) Presence of grooming modulation does not contribute to genotype differences (genotype: p < 0.001; groom modulation: p = 0.01; genotype x groom mod: p = 0.56). However, cells inhibited by grooming in both genotypes are modulated in response to correct lever press more weakly compared with cells not modulated in response to grooming. (f) Similar to (e) but post-fluoxetine treatment (genotype: p = 0.67; groom modulation: p = 0.01; genotype x groom mod: p = 0.34; post-hoc t-test groom inhib vs no modulation: p = 0.001). (g) Similar to Figure 5h, baseline deficits in modulation in response to reward cues normalize after fluoxetine in KOs (genotype: p < 0.001; drug: p = 0.45; genotype x drug: p < 0.001). (h) Prior to fluoxetine treatment, cells inhibited by grooming in both genotypes were modulated in response to reward cues more weakly compared with cells not modulated in response to grooming. However, presence of grooming modulation does not contribute to genotype differences (genotype: p = 0.01; groom modulation: p = 0.04; genotype x groom mod: p = 0.85). (i) Similar to h but post-fluoxetine treatment (genotype: p = 0.81; groom modulation: p = 0.03; genotype x groom mod: p = 0.73). NA indicates that no groom_exc_ neurons were modulated in response to reward cues. P-values of the repeated measures ANOVA are indicated in the inset statistics such that, for panels (d,g), G: genotype, D: drug, GxD: interaction. For panels (e,f,h,I), G: genotype, Groom: grooming modulation, Interaction: genotype x grooming modulation interaction. Asterisks indicate post-hoc t-tests with p-values that are less than 0.05.

**Figure 8:**
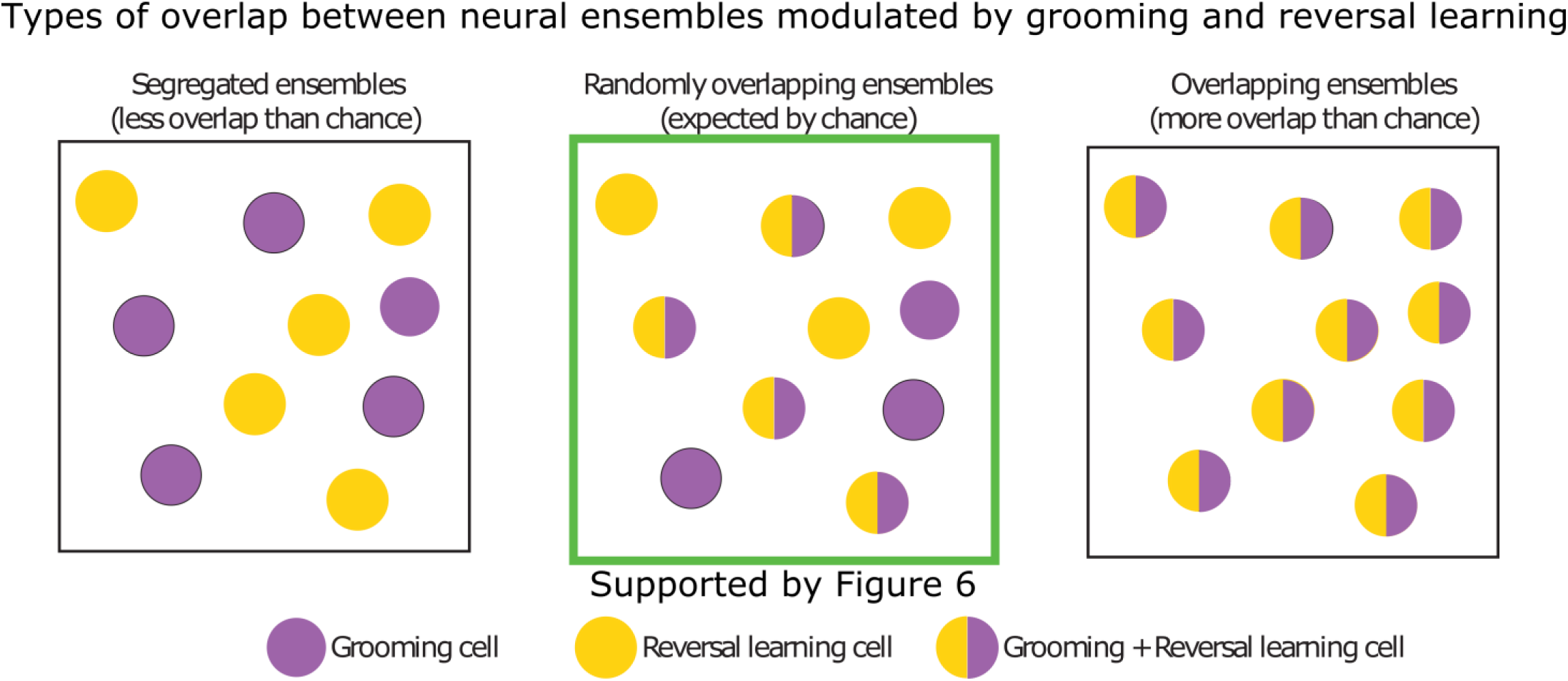
Summary of experimental evidence demonstrating that modulation of compulsive grooming and reversal learning is independent. Grooming and reversal learning could overlap in 3 ways. Grooming and reversal learning cells could be part of separate (left), randomly overlapping (middle) or preferentially overlapping (right) ensembles. Data from Figure 6 suggests that grooming and reversal learning cells are randomly overlapping and are therefore distributed independently of one another.

## Discussion

Patients with OCD display disrupted performance during reversal learning tasks, yet whether these behaviors share common neural underpinnings is unknown. Here we replicate and extend prior work showing that *Sapap3*-KO mice display both compulsive behavior and deficits in reversal learning. While we demonstrate that both deficits improve with fluoxetine, the severity of compulsive grooming does not correlate with the extent of reversal learning deficits. Additionally, we provide several pieces of new evidence suggesting that reversal learning deficits and compulsive behaviors arise from disparate network abnormalities in the LOFC. First, compulsive grooming and deficits in reversal learning are associated with distinct patterns of abnormal LOFC activity, both of which normalize after fluoxetine treatment. During grooming, KO mice have a higher percentage of inhibited LOFC neurons, while during reversal learning, KO mice show weaker modulation in response to the correct lever press and reward cue, with no change in percentages of modulated cells. Second, these patterns of abnormal LOFC activity are independent of one another. We find that reversal learning-modulated neurons are distributed randomly among grooming-modulated neurons. Additionally, the presence of grooming-associated modulation does not contribute to the weaker modulation in response to reversal learning observed in KO mice. Taken together, these data indicate that the LOFC plays both independent and distinct roles in these two pathological behaviors and their improvement with fluoxetine.

Our *in vivo* calcium imaging data suggest that LOFC neurons display heterogeneous excitatory and inhibitory responses to normal grooming in WT mice, and that the specific changes in activity that contribute to pathologic grooming arise from a preponderance of groom_inh_ neurons in KOs. Additionally, our data showing that reductions in grooming are correlated with fluoxetine-associated decreases in groom_inh_ neurons, support an emerging model in which interventions that boost net LOFC activity in *Sapap3*-KOs lead to decreases in grooming (15). How might these subpopulations of groom_inh_ neurons arise? We propose that decreases in activity in groom_inh_ neurons are most likely due to decreases in the excitatory drive (as opposed to an increase in inhibitory drive to groom_inh_ neurons), as post-mortem data from patients with OCD show decreases in a variety of transcripts associated with excitatory, but not inhibitory synapses, including *SAPAP3* (23). It is also possible that these decreases in excitation stem from sources extrinsic to the LOFC. Given that patients with OCD show hyperactivity in cortical-striatal-thalamo-cortical circuits (11–13), at first glance it seems likely that LOFC neurons receive increased, not decreased, inputs from the thalamus, suggesting that reductions in input may arise from other cortical or subcortical areas such as the amygdala. However, we cannot rule out the possibility that groom_inh_ neurons receive increased inhibition during grooming from local interneurons (26). Future studies measuring grooming-related activity selectively in GABAergic interneurons or excitatory neurons that receive inputs from specific areas will be necessary to understand the drivers of the heterogeneous grooming-related activity that we find here.

By taking advantage of the ability to track the same neurons over time, we also began to elucidate how fluoxetine changes how LOFC neurons are modulated in response to behavior. Neurons can belong to one of three populations with respect to modulation in response to grooming after fluoxetine: they can remain modulated; they can stop being modulated; and they can become newly modulated. We find that fluoxetine specifically decreases the percentage of new groom_inh_ without changing the modulation strength, indicating that neurons that stop being inhibited and neurons that become newly inhibited have similar modulation strengths. These data also suggest that subpopulations of LOFC neurons respond differentially to SSRI administration. Because serotonin is increased broadly following systemic fluoxetine administration, it remains unknown whether differential effects of serotonin are due to local effects on the excitability of LOFC subpopulations or whether serotonin modulates other brain areas that preferentially project to subsets of LOFC neurons. As a caveat, we are unable to address the stability of grooming modulation in the absence of fluoxetine or the time frame by which new neurons are added and old neurons drop out of the grooming ensemble as we only analyzed activity at two time points: pre- and post-fluoxetine.

In addition to assessing compulsive grooming, we explored the neural correlates of reversal learning, and found that fluoxetine-associated increases in the strength of modulation in response to correct lever presses and the reward cue by individual LOFC neurons in KO mice parallel improvements in baseline reversal learning deficits. One benefit of our task design is that mice are trained on a VR2 schedule, which allows for the dissociation of neurons associated with the action of the lever press from the sensory cues associated with the reward. As a result, deficits in modulation in response to reversal learning may arise due to disruptions with associating the correct lever press with the reward cue. While the necessity of the LOFC in reversal learning tasks is controversial (35, 36), it is likely involved in mediating the impact of serotonin on reversal learning, as depletion of serotonin in the LOFC is sufficient to disrupt reversal learning (37). While we did not directly assess the necessity of the LOFC for fluoxetine-associated improvements, our calcium imaging data suggest that the LOFC is involved in the normalization of behavior after fluoxetine. Additionally, we cannot assess how value is represented in LOFC neurons given the deterministic nature of the current design, or whether neurons representing correct/incorrect lever presses are modulated in response to that specific lever or rewarded/unrewarded actions in general. Finally, our current encoding model is unable to detect neurons that become modulated, or stop being modulated, within a session. Future task designs in which rewards are probabilistic and contingencies switch multiple times during a single session will be better able to address these limitations.

If disrupted reversal learning and compulsive behaviors arise from distinct and independent coding deficits in the LOFC, what can we conclude about the association between abnormal cognitive flexibility and OCD? Our data may help to reconcile competing interpretations of prior work showing that both patients with OCD and their unaffected relatives have abnormal OFC activity during reversal learning (10). One interpretation is that abnormal OFC activity during reversal learning may constitute an OCD endophenotype and provide a causal and quantifiable link between genetic susceptibility and the symptoms of OCD (38). A second is that, because abnormal OFC activity of unaffected relatives is dissociable from compulsive behaviors, deficits in reversal learning are separable from compulsive behaviors. In this interpretation, OCD risk genes cause a variety of molecular and circuit-level changes: one subset of changes leads to the symptoms of OCD while a separate subset leads to deficits in reversal learning that may impact functioning independent of obsessions and compulsions. The data we report here support the second interpretation – that deficits in reversal learning are unrelated to compulsive behaviors – since they arise from distinct and independent coding deficits.

## Acknowledgments / Disclosures

We would like to thank Xiaojun Li, Brittany Chamberlain and Alexander Lammers for help with video scoring; Drs Jesse Wood, Zoe LaPalombara and James Hyde for help with data processing pipeline development; and the rest of the Ahmari lab for helpful feedback. This work was supported by NIMH R21 MH116330, Burroughs Wellcome CAMS Award, NIMH BRAINS R01MH104255, McKnight Scholar Award, MQ Fellows Award, and Klingenstein-Simons Fellowship Award in the Neurosciences to SEA.

This the article has been posted on BioRxiv preprint server. All authors declare no disclosures or conflicts.

## Supplemental Figures

**Figure S1:**
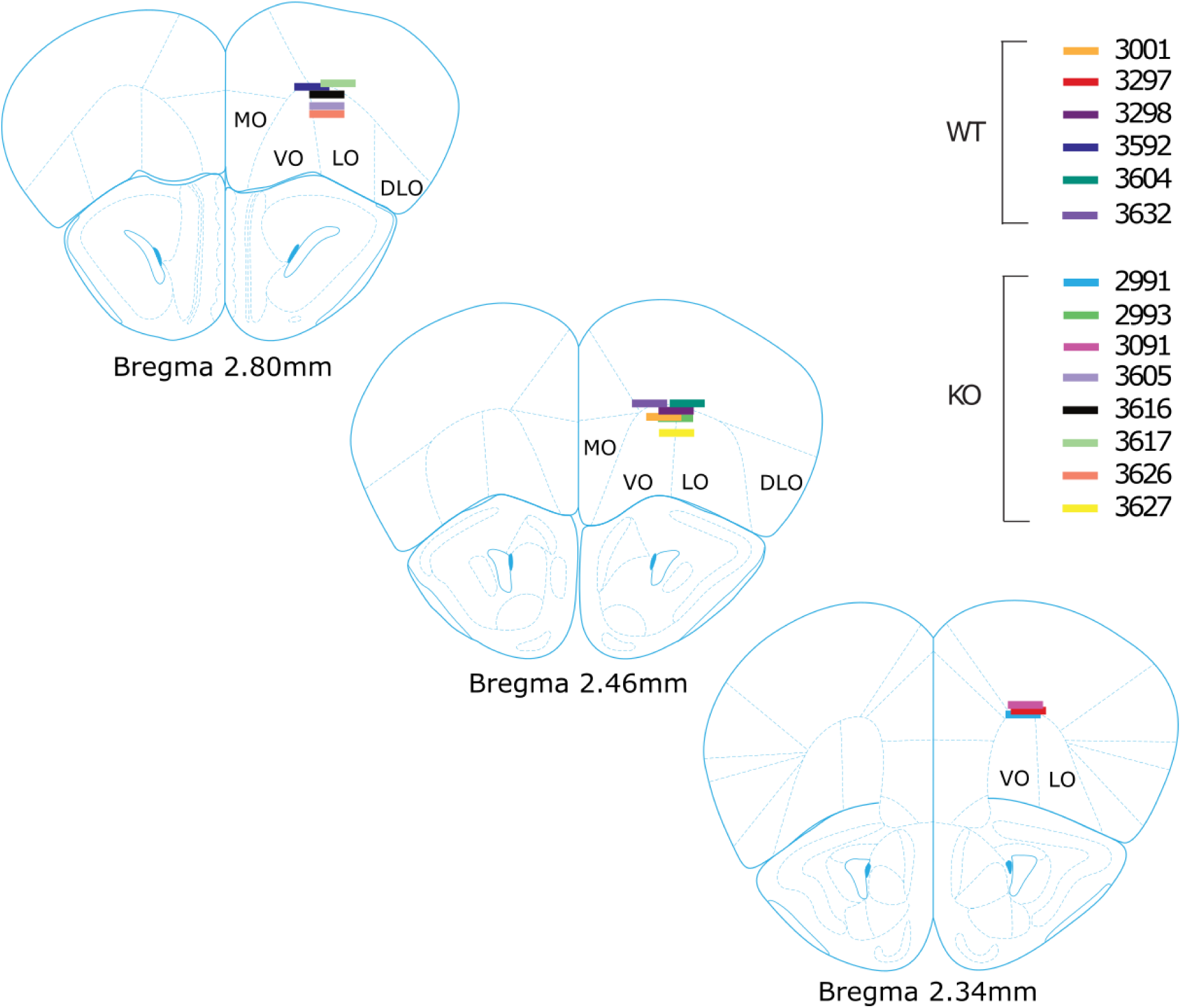
**LOFC lens placements across cohort**

**Figure S2:**
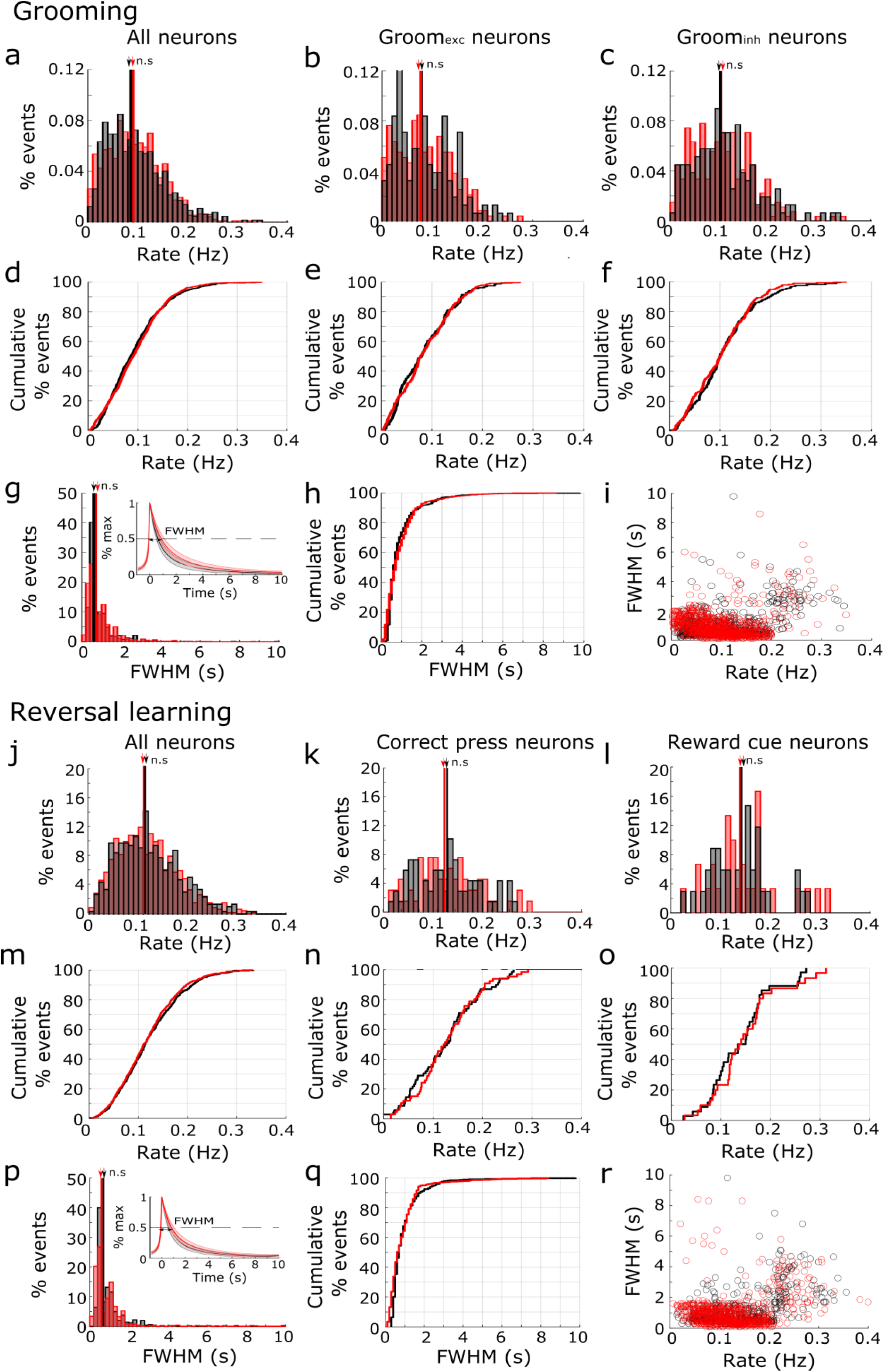
Over 90% of imaged neurons have calcium event properties characteristic of excitatory neurons. To determine whether distinct clusters of neurons were present in our data, event rates and event widths were measured for all cells recorded during the (a-i) pre-fluoxetine grooming session and the (j-r) pre-fluoxetine reversal learning session. Pre-fluoxetine grooming session: (a-c) Histograms of event rates across (a) all neurons, (b) groom_exc_ neurons, and (c) groom_inh_ neurons for WT (black) and KO (red) cells. Median event rates (black and red arrows) were not significantly different between WT and KO cells. (d-f) Same as a-c but cumulative distribution function (CDF) plots. Over 90% of cells had rates less than 0.2 Hz. (g) Histogram of event widths (FWHM=full width at half max) in seconds across cells from WT and KO animals. *Inset:* Average event trace across all events and cells from WT (black) and KO (red) animals. (h) CDF of event widths. Over 90% of cells had event widths less than 2s. i) Plot of event rate vs event width shows a distinct cluster of ∼10% of neurons that have a larger width and faster rate compared with the majority of neurons. Pre-fluoxetine reversal learning session: (j-l) Histograms of event rates across (j) all neurons, (k) neurons modulated in response to correct lever presses, and (l) neurons modulated in response to reward cues for WT (black) and KO (red) cells. Median event widths (black and red arrows) were not significantly different between WT and KO cells. (m-o) Same as j-l but CDF plots. Over 90% of cells had rates less than 0.2 Hz. (p) Histogram of event widths (full width at half max) in seconds across cells from WT and KO animals. *Inset:* Average event trace across all events and cells from WT (black) and KO (red) animals. (q) CDF of event widths. Over 90% of cells had event widths less than 2s. r) Plot of event rate vs event width shows a distinct cluster of ∼10% of neurons that is similar to the grooming dataset. Event times used to calculate event rate and width are identified by CNMFe. Event width is calculated by first averaging across all events from each neuron and then calculating the full width at half-max of the mean event trace.

**Figure S3:**
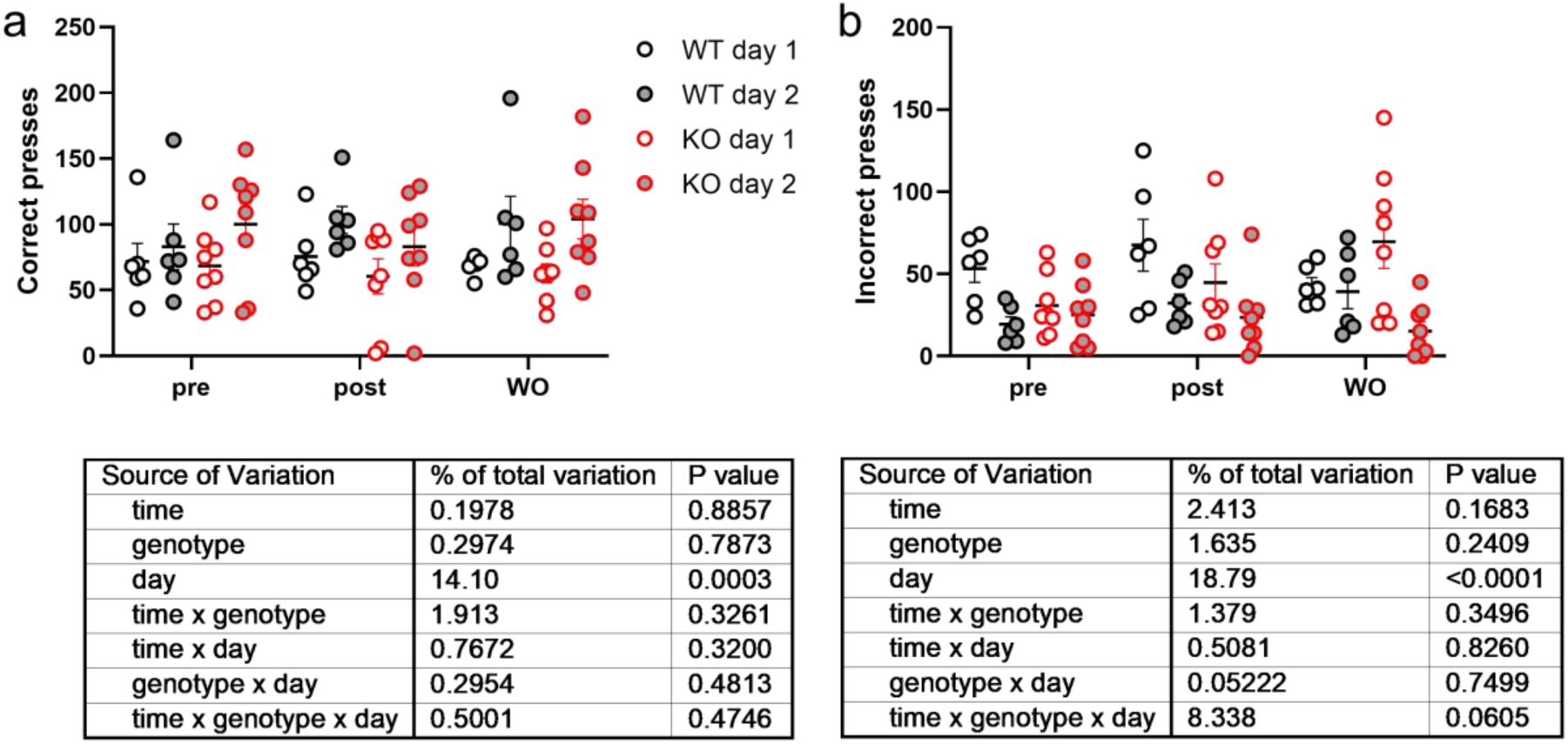
No genotype differences in pre-reversal discrimination were observed. (a) *Top:* There were no differences in correct lever presses between WTs (black) and KOs (red) at either the first (open circles) or second (filled circles) training day for the pre-fluoxetine, post-fluoxetine or washout time points. *Bottom:* Statistical tests for incorrect lever presses. (b) Same as (a) but for incorrect lever presses.

**Figure S4:**
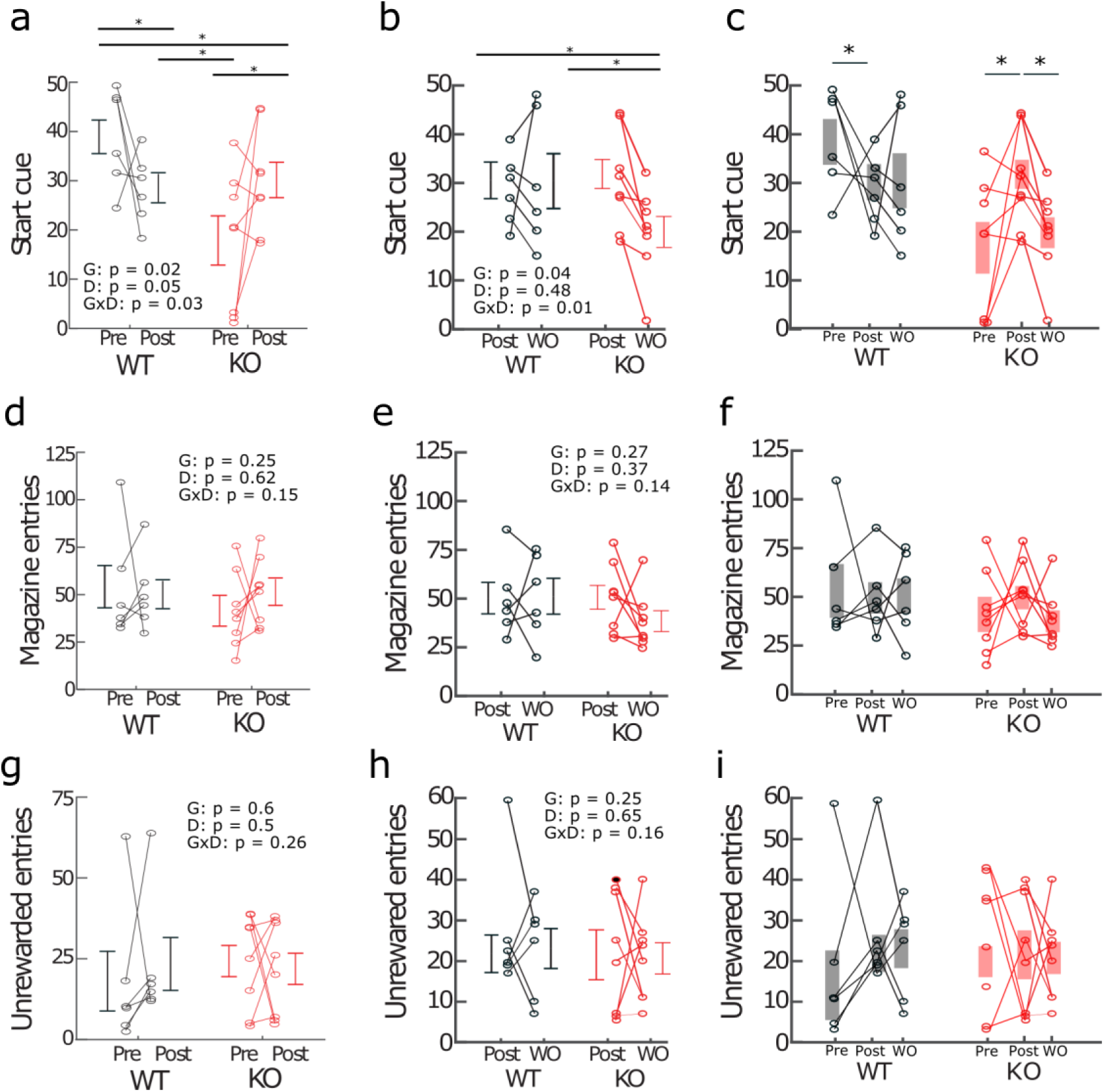
Additional behavioral data from reversal learning task. (a) Trial start cues pre- and post-fluoxetine (genotype: F_(1,12)_ = 6.9, p = 0.02; drug: F_(1,12)_ = 0.05, p = 0.83; genotype x drug: F_(1,12)_ = 6.6, p = 0.025). (b) Trial start cues post-fluoxetine and after washout (genotype: F_(1,12)_ = 5.38, p = 0.04; drug: F_(1,12)_ = 0.48, p = 0.51; genotype x drug: F_(1,12)_ = 8.0, p = 0.01). (c) Trial start cues plotted pre-fluoxetine, post-fluoxetine and after washout. Data from a-b. (d) Total magazine entries pre- and post-fluoxetine (genotype: F_(1,12)_ = 1.45, p = 0.25; drug: F_(1,12)_ = 0.25, p = 0.62; genotype x drug: F_(1,12)_ = 2.31, p = 0.15). (e) Total magazine entries post-fluoxetine and after washout (genotype: F_(1,12)_ = 1.32, p = 0.27; drug: F_(1,12)_ = 0.86, p = 0.37; genotype x drug: F_(1,12)_ = 2.51, p = 0.14). (f) Total magazine entries plotted pre-fluoxetine, post-fluoxetine and after washout. Data from d-e. (g) Unrewarded magazine entries pre- and post-fluoxetine (genotype: F_(1,12)_ = 0.29, p = 0.6; drug: F_(1,12)_ = 0.49, p = 0.5; genotype x drug: F_(1,12)_ = 1.34, p =0.26). (h) Unrewarded magazine entries post-fluoxetine and after washout (genotype: F_(1,12)_ = 1.45, p = 0.25; drug: F_(1,12)_ = 0.21, p = 0.65; genotype x drug: F_(1,12)_ = 2.29, p = 0.16). (i) Unrewarded magazine entries plotted pre-fluoxetine, post-fluoxetine and after washout. Data from g-h. Asterisks indicate post-hoc t-tests with p-values that are less than 0.05.

**Figure S5:**
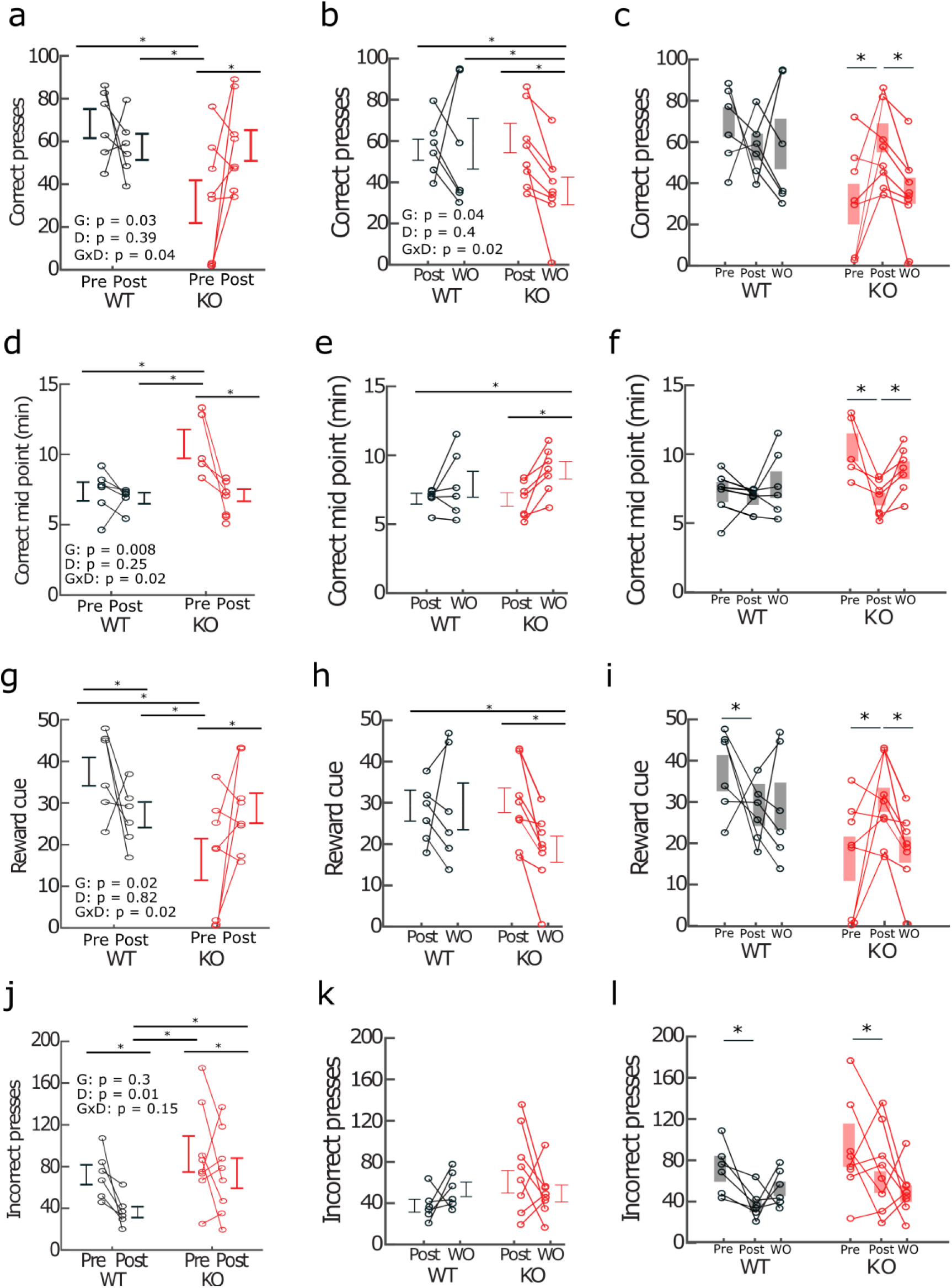
Improvements in reversal learning after fluoxetine treatment disappear after washout (WO). (a) Correct lever presses pre- and post-fluoxetine. Same data as Figure 4c. (b) Correct presses post-fluoxetine and after washout (genotype: F_(1,12)_ = 5.4, p = 0.04; drug: F_(1,12)_ = 0.76, p = 0.4; genotype x drug: F_(1,12)_ = 6.78, p = 0.02). (c) Correct lever presses plotted pre-fluoxetine, post-fluoxetine and after washout. Data from a-b. (d) Time to acquire correct lever presses pre- and-post fluoxetine. Data from Figure 4e. (e) Time to acquire correct presses post-fluoxetine and after washout (genotype: F_(1,12)_ = 4.7, p = 0.051; drug: F_(1,12)_ = 0.04, p = 0.81; genotype x drug: F_(1,12)_ = 6.5, p = 0.03). (f) Time to acquire correct lever presses plotted pre-fluoxetine, post-fluoxetine and after washout. Data from d-e. (g) Reward cues pre- and post-fluoxetine. Same data as Figure 4f. (h) Reward cues post-fluoxetine and after washout (genotype: F_(1,12)_ = 6.82, p = 0.02; drug: F_(1,12)_ = 0.04, p = 0.83; genotype x drug: F_(1,12)_ = 6.61, p = 0.02). (i) Reward cues plotted pre-fluoxetine, post-fluoxetine and after washout. Data from g-h. (j) Incorrect presses pre- and post-fluoxetine. Same data as Figure 4g. (k) Incorrect presses plotted post-fluoxetine and after washout (genotype: F_(1,12)_ = 0.78, p = 0.38; drug: F_(1,12)_ = 0.1.3, p = 0.26; genotype x drug: F_(1,12)_ = 2.31, p = 0.15). (l) Incorrect presses plotted pre-fluoxetine, post-fluoxetine and after washout. Data from j-k. Asterisks indicate post-hoc t-tests with p-values that are less than 0.05.

**Figure S6:**
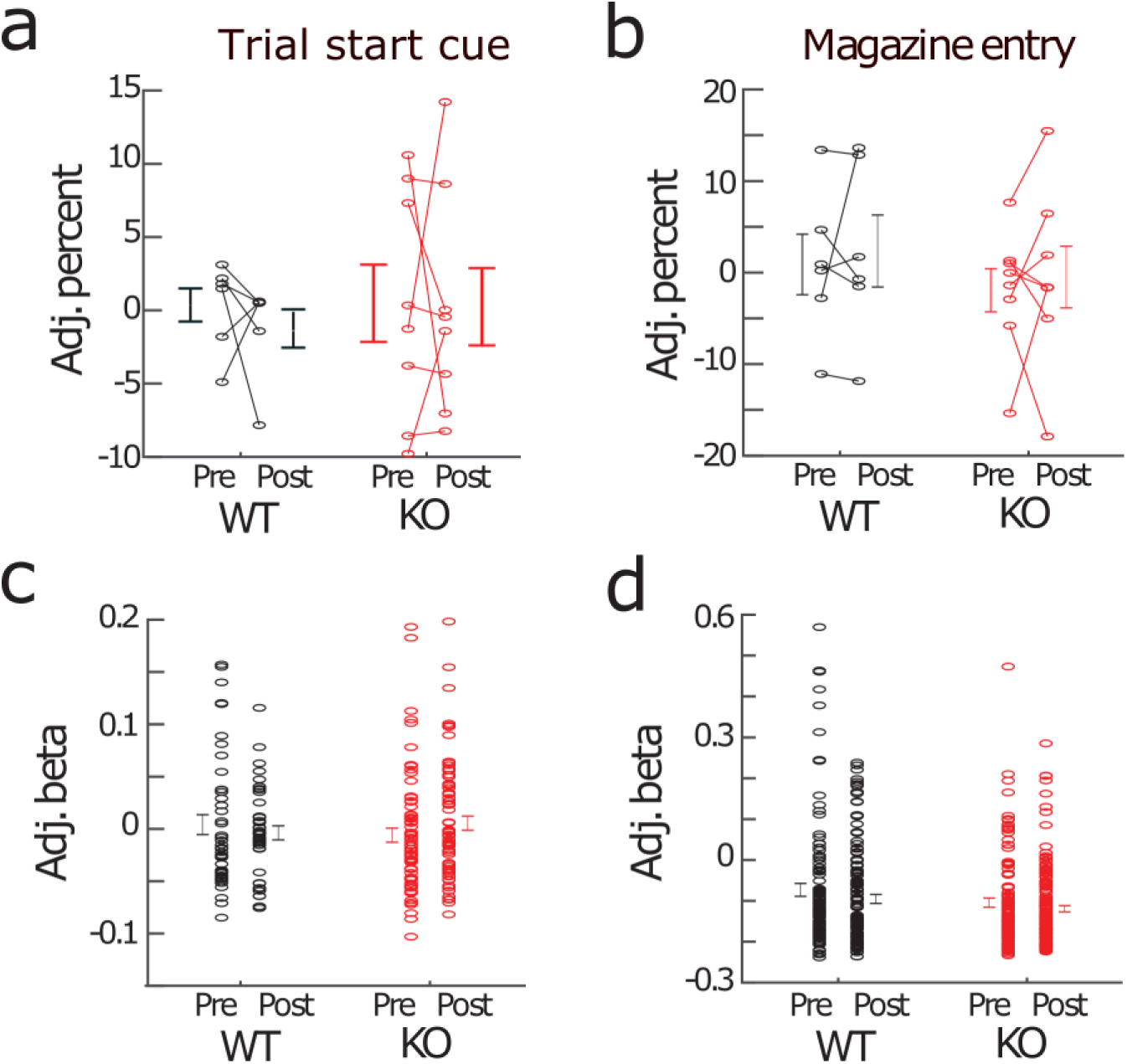
Additional imaging data from the reversal learning task. (a-b) Genotype and fluoxetine effects on the percentage of cells modulated in response to (a) trial start cue (genotype: F_(1,12)_ = 1.48, p = 0.24; drug: F_(1,12)_ = 0.29, p = 0.6; genotype x drug: F_(1,12)_ = 1.34, p = 0.26) and (b) magazine entry (genotype: F_(1,12)_ = 0.49, p = 0.49; drug: F_(1,12)_ = 0.42, p = 0.52; genotype x drug: F_(1,12)_ = 1×10^-5^, p = 0.99). Variables adjusted for number of task events (adj). (c-d) Genotype and fluoxetine effects on the strength of modulation in response to (c) trial start cue (genotype: p = 0.34; drug: p = 0.54; genotype x drug: p = 0.21) and (d) magazine entry (genotype: p = 0.65; drug: p = 0.29; genotype x drug: p = 0.37. Variables adjusted for number of task events and animal-to-animal variability.

**Figure S7:**
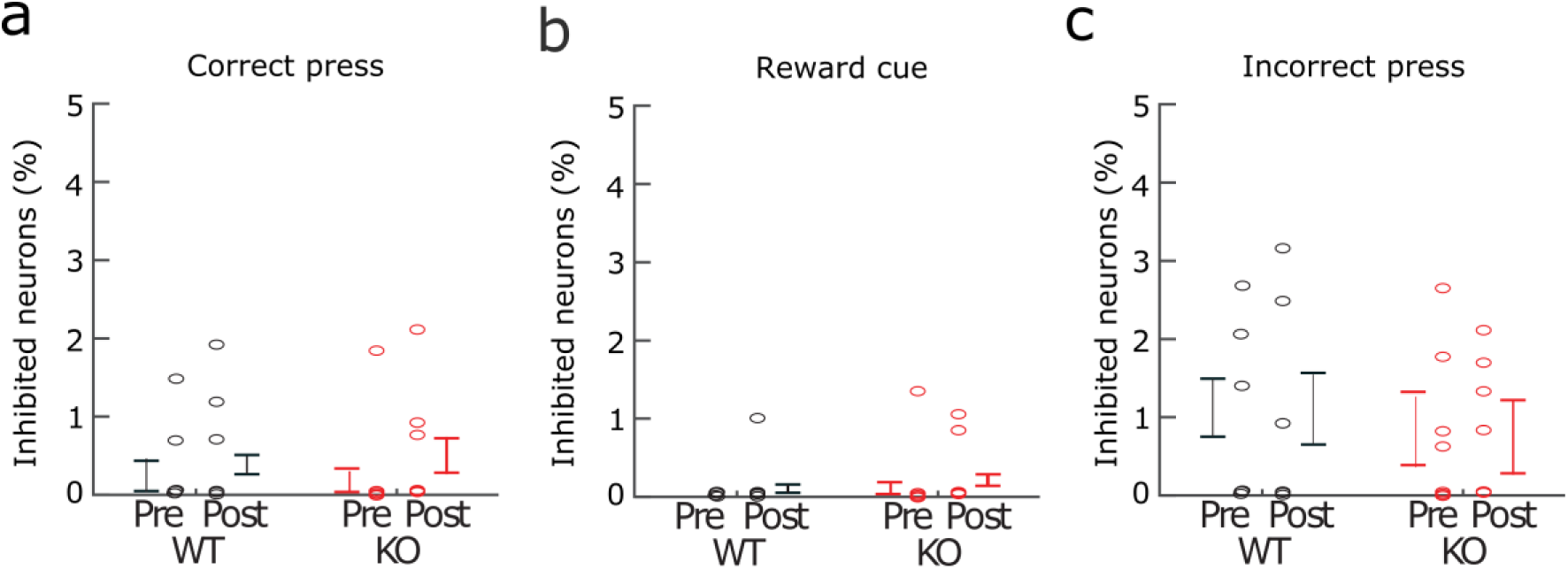
Few neurons were inhibited during reversal learning task events. (a) Percent of neurons inhibited during correct lever presses. (b) Percent of neurons inhibited during reward cue. (c) Percent of neurons inhibited during incorrect lever presses.

**Figure S8:**
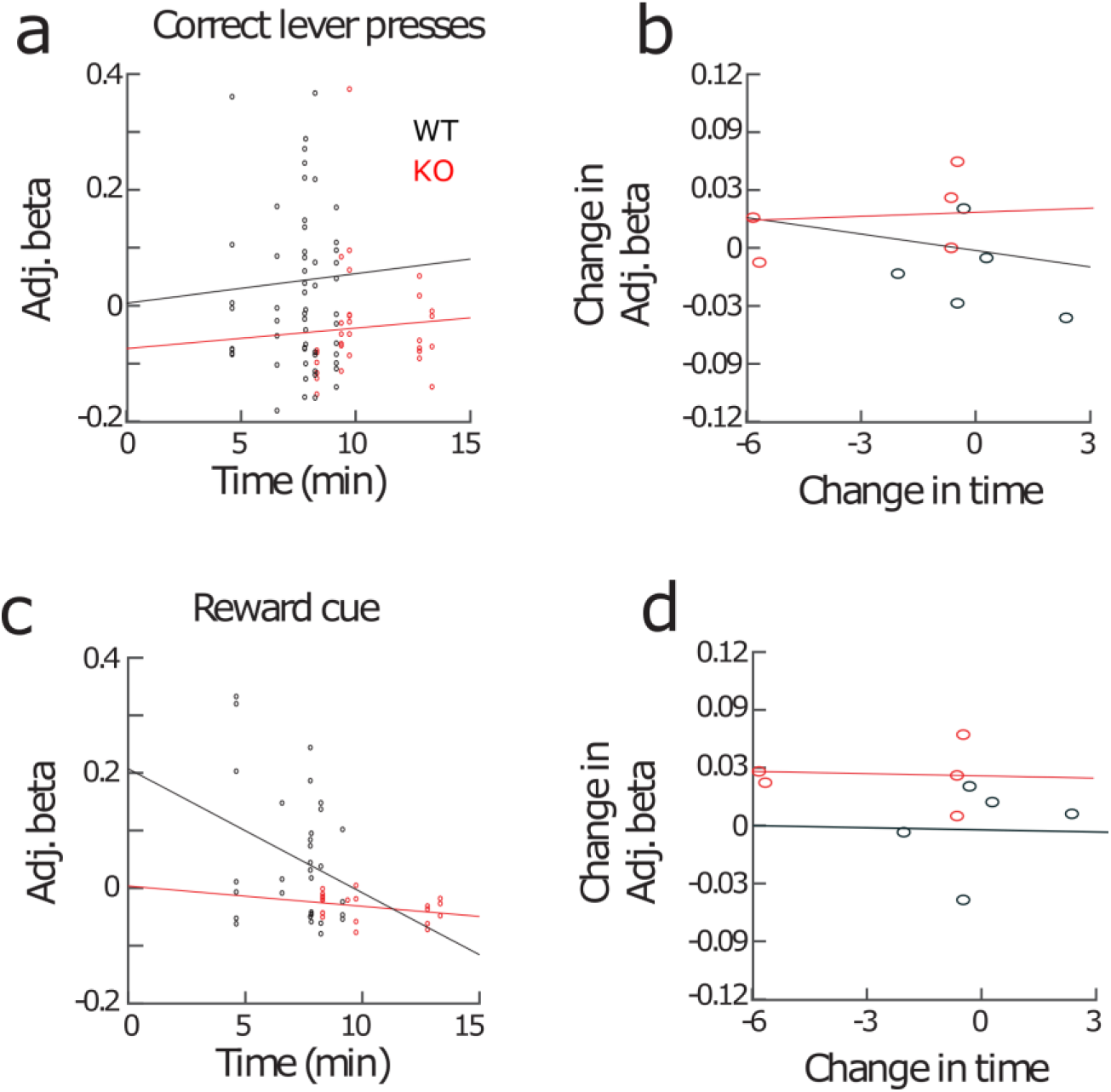
Correlation between strength of modulation and correct midpoint. (a) Adjusted beta weight for correct lever presses vs correct midpoint. (b) Change in adjusted beta weight for correct lever presses pre- and post-fluoxetine vs change in correct midpoint pre- and post-fluoxetine. (c-d) Same as (a-b) but for strength of modulation in response to reward cue. All correlations were not significant.

**Figure S9:**
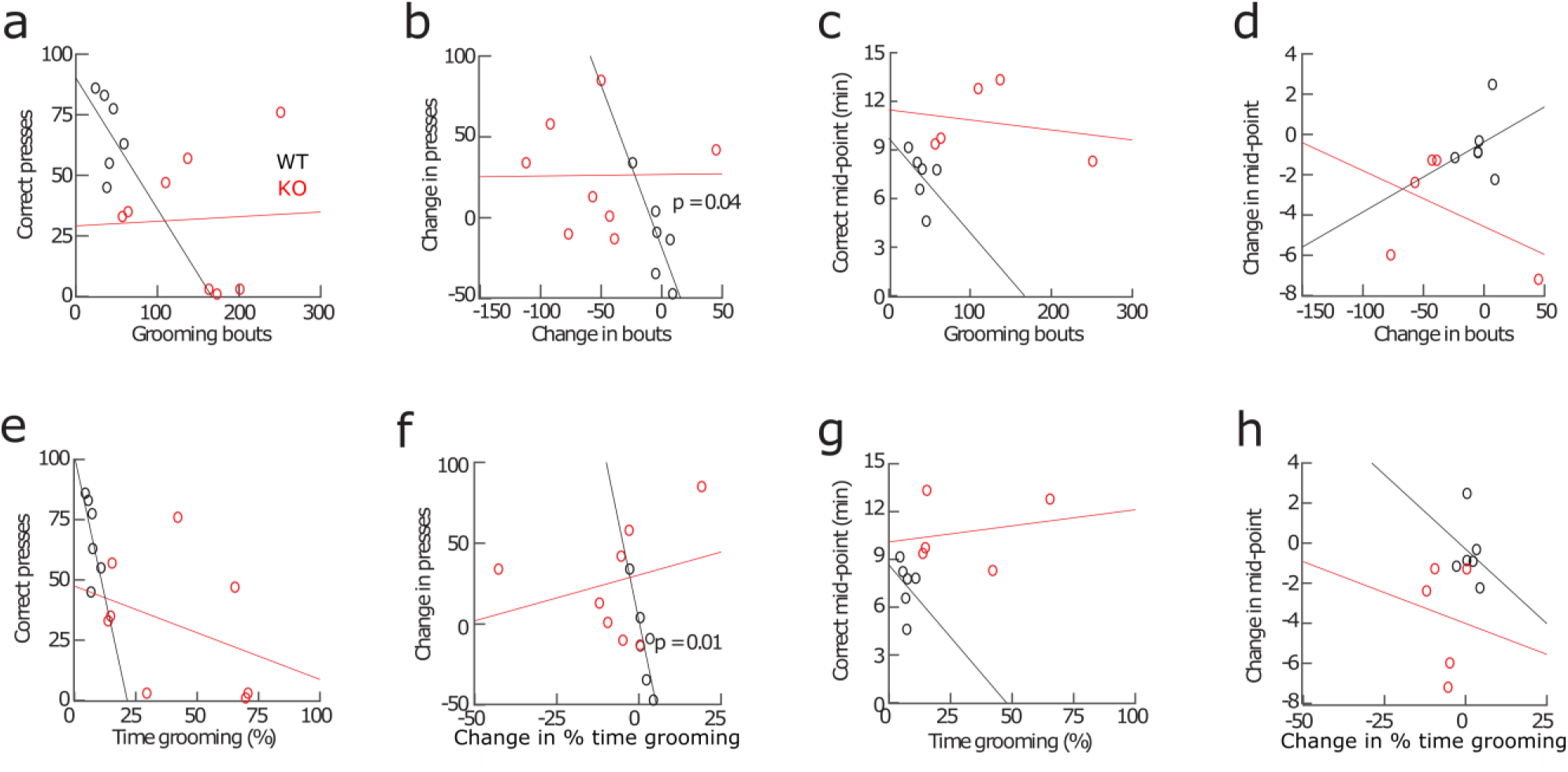
Correlations between grooming and reversal learning behavior. (a) Correct lever presses versus grooming bouts. (b) Change in correct lever presses versus change in grooming bouts. (c) Mid-point of correct lever presses versus grooming bouts. (d) Change in mid-point versus change in grooming bouts. (e) Correct lever presses versus % time grooming. (f) Change in correct lever presses versus change in % time grooming. (g) Mid-point of correct lever presses versus % time grooming. (h) Change in mid-point versus change in % time grooming. No correlations were significant unless marked on the graphs.

**Figure S10:**
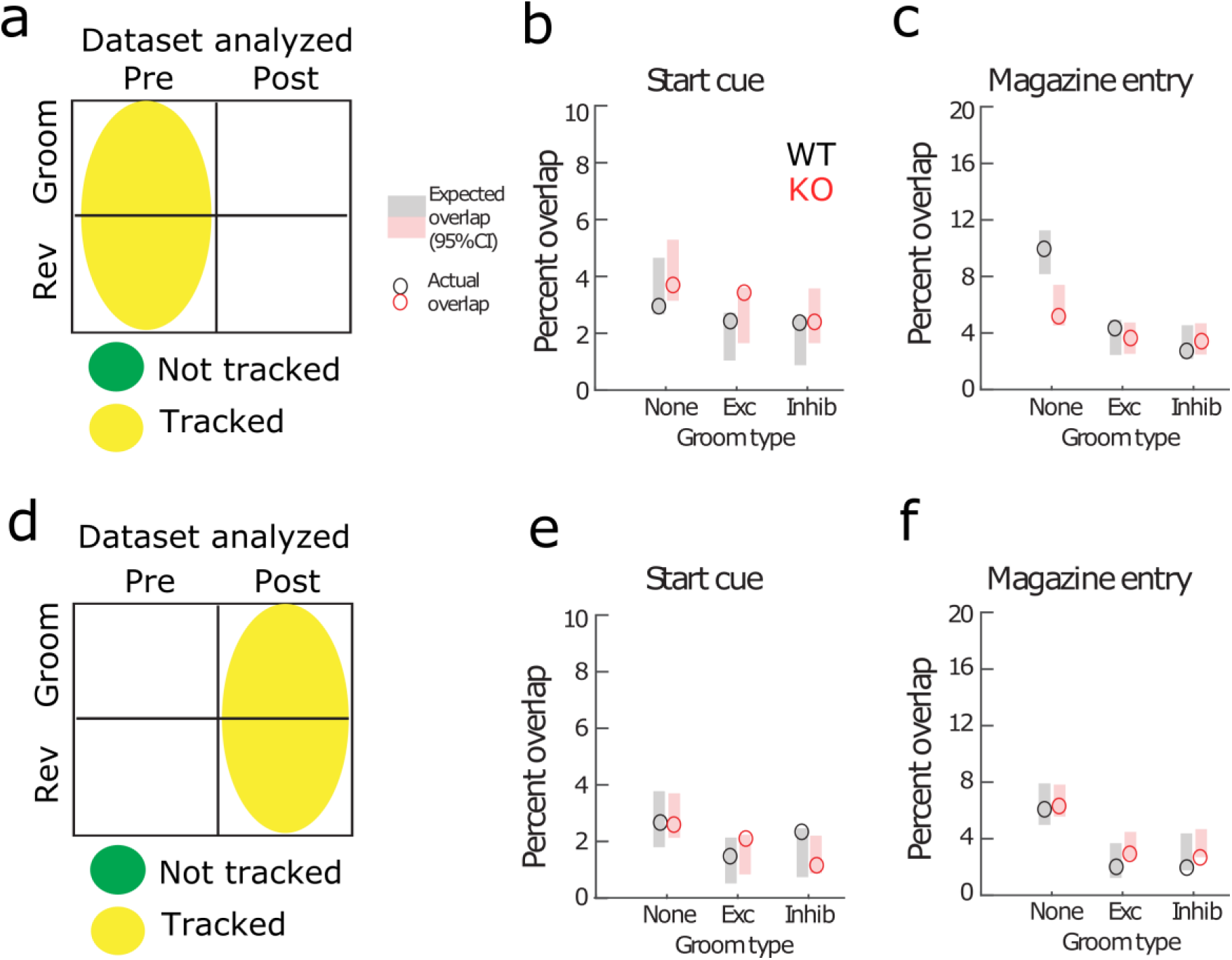
Overlap between additional reversal learning events and grooming. Reversal learning cells are randomly distributed among grooming cells both pre-fluoxetine (a-c) and post-fluoxetine (d-f). Expected overlap (95% CI) between reversal learning and grooming cells (shaded rectangles). Actual overlap between reversal learning and grooming cells (circles). (b) Trial start cue. (c) Magazine entries. (d-f) Same as (a-c) but for cells tracked between post-fluoxetine grooming and reversal sessions.

**Figure S11:**
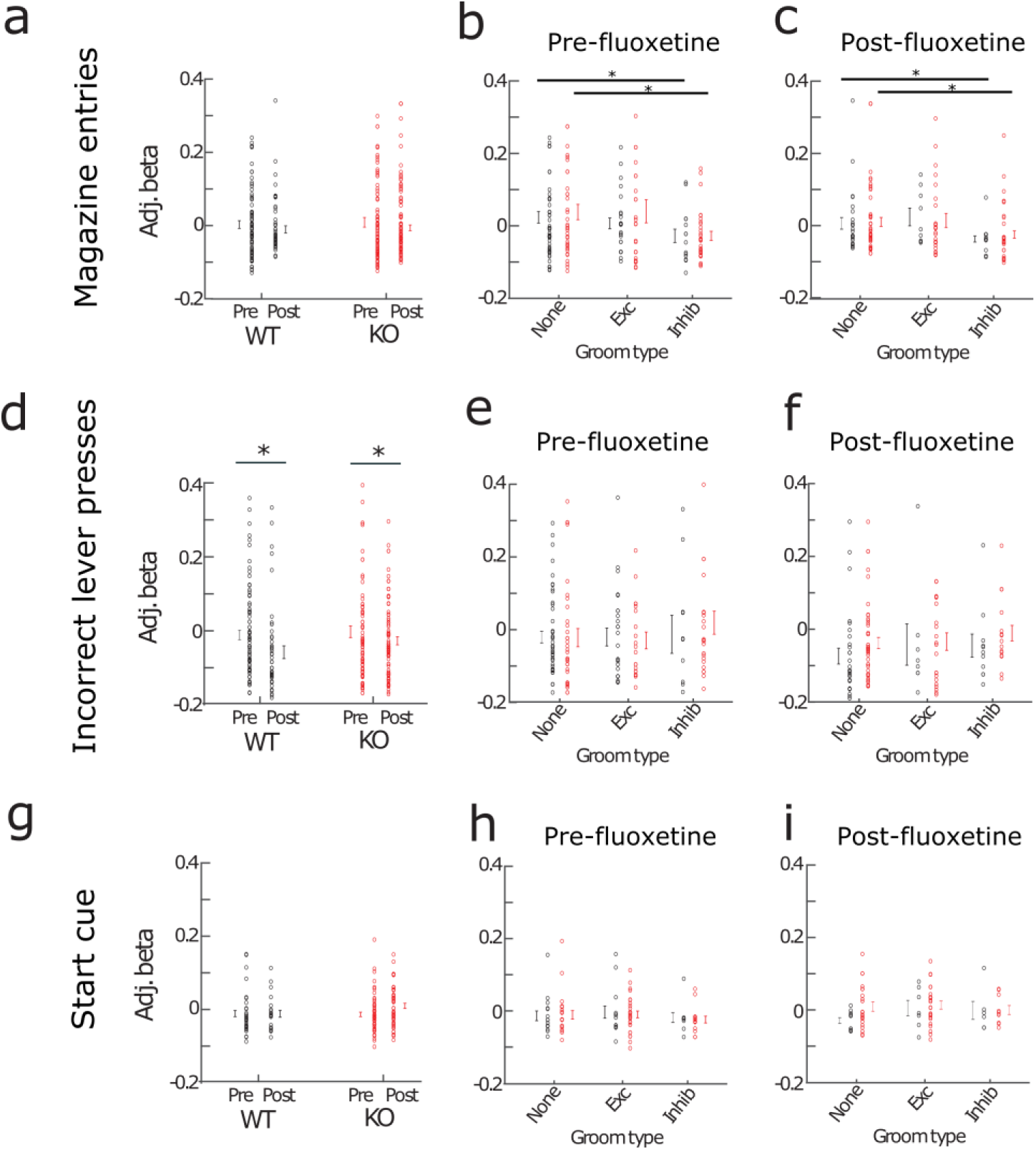
Interaction between additional reversal learning events and grooming. Cells are tracked between grooming and reversal sessions at both the pre- and post-fluoxetine time points. Cells are not tracked between the pre- and post-sessions. (a) Strength of modulation in response to magazine entries pre- and post-fluoxetine. (b) Strength of modulation in response to magazine entries pre-fluoxetine grouped by how the cell is modulated in response to grooming (genotype: p = 0.48; groom modulation: p = 0.02; genotype x groom mod: p = 0.28). (c) Similar to (b) but post-fluoxetine treatment (genotype: p = 0.43; groom modulation: p = 0.01; genotype x groom mod: p = 0.61). (d-f) Same as (a-c) but for incorrect lever presses. (d) Strength of modulation pre- and post-fluoxetine (genotype: p = 0.56; drug: p = 0.02; genotype x drug: p = 0.21). (g-i) Same as (a-c) but for trial start cue.

## Methods supplement

### Animals

*Sapap3*-knockout (*Sapap3*-KO) and wildtype (WT) littermates were generated through breeding *Sapap3* heterozygous mutants (*Sapap3*^+/-^). *Sapap3*-KO (n = 8; 5 female) and WT (n = 6; 3 female) littermates were 6-7 months old at the time of first surgery, and imaging experiments were conducted starting at 9-10 months of age to ensure robust expression of compulsive grooming phenotype in *Sapap3*- KOs (1).

### Calcium imaging surgery

Mice were anesthetized using 5% isoflurane mixed with oxygen and maintained on 1-2% isoflurane for the duration of surgery. Mice were placed on a small-animal stereotactic instrument (Kopf Instruments, Tujunga, CA) and secured using ear bars and a bite bar. Hair was removed from the dorsal surface of the head with hair clippers and the incision area was scrubbed with a betadine solution. A large incision was then made exposing the dorsal portion of the skull. AP and ML measurements were made relative to an interpolated bregma; DV measurements were made relative to dura. 800nl of a virus encoding GCaMP6f under the synapsin promotor (AAV5-synapsin-GCaMP6f-WPRE-SV40, titer 1.82×10^12^; Penn Vector Core) was injected into the LOFC adjacent to the lens implant target (AP: +2.7, ML: -1.0, DV: - 1.6) using a fixed needle Hamilton syringe (Cole-Parmer Scientific, Vernon Hills IL, USA) connected to sterile polyethylene tubing affixed to a metal cannula and a Harvard Apparatus Pump 11 Elite Syringe Pump (Harvard Apparatus, Holliston MA, USA). Immediately after injection of virus into LOFC, a 500µm diameter, 6.1mm length gradient refractive index lens (ProView GRIN lens, Inscopix Pala Alto, CA USA) was lowered just dorsal to the viral injection target (AP: +2.6, ML: -1.2, DV: -1.4) to allow for visualization of cells in the target region. GRIN lenses were secured in place with black dental cement (Ortho-Jet, Lang Dental, Wheeling, IL) surrounding the lens and two 0.45mm skull screws were placed just anterior of the lambdoid suture. Following completion of each surgery, mice were injected with subcutaneous (s.c.) carprofen (3mg/kg in 0.9% saline; Henry Schein, Melville, NY) and administered topical antibiotic ointment (TAO; Henry Schein) and lidocaine HCL 2% ointment (Henry Schein) around the headcap. Mice were then placed on a heating pad and given DietGel (ClearH_2_O, Portland ME, USA) and monitored until they were fully recovered from anesthesia. Mice were administered carprofen s.c. and received lidocaine and TAO treatments for 3 days post-surgery. For all surgical procedures mice were kept group housed with same sex littermates.

After ∼3-4 weeks for viral incubation, a second procedure was performed during which mice were again anesthetized with isoflurane and secured to a stereotactic apparatus using ear cuffs. Using a Dremel, excess dental cement was carefully removed exposing the ProView GRIN lens. The top of the ProView lens was then cleaned with compressed air, lens paper, and 100% ethanol to remove all dental cement dust. A magnetic microscope baseplate (Part ID:1050-002192, Inscopix) was then attached to the miniaturized microscope (nVistaHD 2.0 epifluorescence microscope, Inscopix) and lowered into place above the GRIN lens with the 475nm blue LED gain and power increased until the lens and gross structures were visible. With isoflurane maintained at 0.5-1%, the optimal field of view was then determined by focusing on visible/active cells or other gross landmarks (blood vessels). Once an optimal field of view was obtained, the baseplate was cemented in place and a plastic Microscope Baseplate Cover (Part ID:1050-002193; Inscopix) was attached to prevent debris from blocking the lens.

### Fluoxetine preparation and administration

(±)Fluoxetine hydrochloride (Fluoxetine; NIMH Chemical Synthesis and Drug Supply Program) was administered via drinking water according to established methods (2, 3). Briefly, bottles were placed in each cage and drinking was monitored for 3 consecutive days to calculate the concentration necessary to achieve the target 18 mg/kg dose, which produces serum fluoxetine levels comparable to a high dose of fluoxetine that is efficacious in OCD patients (3). Based on the average daily consumption and the average weight of the mice in each cage (and in line with historical averages within our lab), 100mg/L fluoxetine hydrochloride was mixed with autoclaved drinking water and stored in black bottles to prevent light degradation. Fluid consumption and bodyweight was continually monitored weekly and bottles were changed every 4 days to avoid degradation of fluoxetine. After 4.5 weeks of administration, fluoxetine was removed from cages and mice were given a two-week washout from the drug to evaluate whether behavior differences returned to baseline levels.

### Experimental timeline

#### Experimental design

Mice underwent testing in a repeated measures design, in which all subjects were treated once with a 4-week dosing regimen of fluoxetine, and within subject data was compared between baseline, fluoxetine treatment, and drug washout periods. Behavior and neural data were collected weekly for grooming and typically every other week for reversal learning (Fig 1b). On weeks where both behaviors were tested, grooming was tested first, food was removed from the cages that afternoon, and mice underwent operant training for 2 days prior to reversal on the third day (see details below). Grooming was tested at baseline, following 1, 2, 3, and 4 weeks of treatment, and at 1 and 2 weeks during drug washout. Reversal was tested at baseline, 1, 3, and 4 weeks treatment, and 2 weeks washout. Data presented here are for baseline and 4 weeks treatment (neural and behavioral data) and washout (reversal behavior data only).

#### Habituation

Following recovery from baseplate surgery, mice underwent extensive habituation using both the miniature nVista 2.0 microscope, and a “model” microscope that is the same size and weight and can be attached to the animal’s baseplate. Mice were extensively habituated in testing environments used for both grooming and operant training with both model and real microscopes prior to these studies.

#### Prior testing

Prior to baseline testing, mice underwent >2 weeks operant training (similar to (1)) for acquisition of lever pressing (fixed ratio 1 schedule, 1 hour sessions), training on rule 1 (e.g. left lever correct, right lever incorrect; counterbalanced, variable ratio (VR) 2 schedule, 30 minute sessions), first reversal (5 days training rule 2) and second reversal (1 day training rule 1). Thus, mice had extensive experience with operant testing and rule changes prior to baseline testing. Mice were also previously tested in at least 2 x 40 minute imaging sessions for grooming analysis similar to procedures used in these studies.

#### Pre-reversal training

Prior to reversal, all mice were trained for 2 consecutive days at each time point on the same rule (e.g. left lever correct) as the last training day during the previous testing period (e.g. correct rule on reversal day from the prior reversal timepoint). No criteria were used to decide whether mice moved onto reversal except completion of these 2 training sessions.

#### Grooming imaging procedures

A custom-built behavioral apparatus was constructed for the accurate simultaneous assessment of grooming behavior and neural activity via *in vivo* calcium imaging. A clear plexiglass sheet was suspended over a behavioral acquisition camera (Point Grey Blackfly, FLIR Integrated Imaging Solutions). A clear acrylic chamber (8”x8”x12”) was placed above the camera such that a mouse could be visualized from below. Behavioral acquisition was conducted at 40 Hz using SpinView (Point Grey, Wilsonville, OR) software and detailed frame information was sent directly to a central data acquisition box (LabJack U3-LV, Labjack Corporation, Lakewood CO USA) which was also receiving calcium frame information (20hz) from nVista software. A pseudorandom flashing (30s ITI) LED visible in the behavioral video controlled by custom scripts via an Arduino (Arduino Leonardo, Somerville MA, USA) and sending TTL pulses to the LabJack was used for alignment of behavior and calcium data. Mice were recorded for 40 minutes during grooming analysis sessions.

Following acquisition, behavioral video was converted and compressed (maintaining accurate frame rate information) into .MP4 format using the open source software HandBrake. Videos were then imported into Noldus The Observer XT (Noldus, Leesburg VA, USA) and grooming behavior was manually scored frame by frame. Grooming behavior was scored as per previous reports (4) by an observer blind to experimental condition (genotype and drug treatment). A mouse was considered to be grooming if it engaged in any of the following behaviors: 1) Facial grooming: mouse touches its face, whiskers, or head with its forepaws. 2) Body grooming: mouse licks its flank or its ventral surface. 3) Hind leg scratching: mouse uses one of its hind legs to scratch its flank/ear/face. The beginning of a grooming bout was defined as the frame when a mouse made a movement to begin grooming (e.g. a face grooming bout began the frame a mouse lifted its paw off the ground to touch its face). The end of a grooming bout was defined as the frame when a mouse ceased grooming (e.g. a body grooming bout ended when a mouse moved its snout from its flank). Grooming bouts separated by less than 500ms were collapsed into a single bout; consequently, the minimum amount of time between grooming bouts for all experiments was 500ms.

#### Operant conditioning imaging procedures

Mice were tested in a reversal learning paradigm using operant chambers containing two retractable levers with cue lights located above the levers. Levers were positioned either side of a reward magazine (Med Associates, Fairfax, VT). For imaging, several modifications were made including: 1) fabrication of custom magazines with Med Associates IR detectors attached which allowed mice to easily access rewards during imaging and allowed detection of head/body entries to the magazine; 2) placement of a textured Perspex floor over the standard bar floor to provide mice with greater stability; 3) extended inner-chamber walls so that the microscope cable could drop straight into the chamber from a hole made in the roof of the sound attenuation chamber, with room for the cable to easily move during exploration of the chamber. MedPC session code included commands to generate outputs corresponding to behavior events (lever press, magazine entry), and these outputs, along with task events of interest (reward delivery, trial start cue), were sent to a central data acquisition box (LabJack) as TTL pulses generated by Med Associates passive connection panel, where they were synchronized in real time with “sync” signals from nVista corresponding to each frame of calcium imaging data acquired.

Mice that had undergone extensive operant training (details in experimental timeline above) were used. For each timepoint mice were tested for three days (30 minutes each). The first two days were tested on the same rule (e.g. left lever correct, right lever incorrect, using the same contingency from their most recent prior test), during which they were trained with the “model” microscope. On the third day the contingency was reversed, and calcium imaging was performed. For all sessions, mice were trained on a VR2 schedule. Upon the start of a trial, both levers were inserted into the chamber and both cue lights above the levers were turned on. When mice completed a rewarded correct response, this resulted in retraction of the two levers, the two cue lights being extinguished, and delivery of a reward pellet (20-mg chocolate-flavored grain-based pellets; BioServ, Flemington, NJ). Reward retrieval (first magazine entry following reward delivery) triggered the start of an inter-trial-interval (ITI) that was randomly set to 5, 6, or 7 seconds duration. If no magazine entry was detected in 10 seconds after reward delivery, the ITI was triggered. At the end of the ITI, a new trial commenced, with the two levers inserted into the chamber and two cue lights turning on. A houselight covered by a red light filter remained lit throughout the session. During studies mice had ad libitum access to food and water, except during operant training when they received restricted food access to maintain 85-90% free feeding body weight. Food was removed on the night prior to operant training, and mice received restricted food access (after operant training) during the 3 day period of operant testing at each timepoint of the study. Neural data and behavior were recorded for 30 minutes but only the first 15 minutes were used for analysis given that learning had occurred within this time period.

#### Calcium imaging acquisition, processing, and analysis

On imaging acquisition days, the microscope was attached and mice were placed into a temporary holding cage. Mice were given 3-5 minutes after attachment of the microscope for recovery from scruffing and to allow any rapid photobleaching to occur. After this period, mice were carefully placed into the testing arena. nVistaHD software recorded compressed greyscale tiff images at 20 Hz. For all mice, analog gain of the image sensor was set between 1 and 4 while the 470 nm LED power was set between 10 to 30% transmission range. A caliper was used to accurately measure the precise microscope focus such that multiple imaging sessions were conducted with the same field of view. These settings were kept consistent for each mouse throughout all subsequent imaging sessions.

All imaging pre-processing was performed using Mosaic software (version 1.2.0, Inscopix) via custom Matlab (MATHWORKS, Natick MA, USA) scripts. Videos were spatially downsampled by a binning factor of 4 (16x spatial downsample) and temporally downsampled by a binning factor of 2 (down to 10 frames per second). Lateral brain motion was corrected using the registration engine TurboReg (5), which uses a single reference frame to match the XY positions of each frame throughout the video. Motion corrected 10 Hz video of raw calcium activity was then saved as a .TIFF and used for cell segmentation.

Using custom Matlab scripts, the motion corrected TIFF video was then processed using the Constrained Non-negative Matrix Factorization approach (CNMFe), which has been optimized to isolate signals from individual putative neurons from microendoscopic imaging (6). Putative neurons were identified and manually sorted by an observer blind to genotype according to previously established criteria (7). After putative neurons were automatically segmented via CNMFe, the spatial footprint and both the raw and denoised calcium traces for each potential neuron were examined by an observer blind to genotype. While blind to genotype, well experienced observers sorted neurons identified via CNMFe. Two main factors were used for determining whether a neuron was included in the analysis. 1) The spatial footprint had to be located within the circular confines of the GRIN lens and have clear and crisp borders characterized by smooth increases in fluorescence from the outside of the footprint to the center. Spatial footprints were typically spherical toward the center of the GRIN lens, and slightly elongated toward the edges due to optical aberrations caused by the lens. 2) The raw and denoised calcium traces had to have detectable phasic calcium transients from above baseline fluorescence (noise) levels.

Custom Matlab (MATHWORKS) scripts were used to align calcium traces with behavior. Grooming behavior (state events) was exported as timestamps (grooming start and grooming stop) and aligned to Ca^2+^ time by recording 5 consecutive LED pulses (point events). The offset of Noldus behavior time to nVista Ca^2+^ time was then subtracted off leaving the same number of frames for both the behavior and Ca^2+^ fluorescence. Grooming timestamps were then transferred to a binary/continuous trace of the same length and sampling rate (10 Hz) as each Ca^2+^ trace via logical indexing (grooming = 1, not-grooming = 0). For operant data alignment, behavior events and calcium imaging frames were logged on the same timescale (by Labjack) in real time, and custom MATLAB scripts were used to extract behavior time stamps relative to calcium imaging session start time. Timestamps for behavior are converted to the closest matching frame in the calcium recording (maximum error of one frame or ± 100ms at 10 Hz).

#### Longitudinal tracking of neurons across sessions

Putative neurons identified via CNMFe were matched using a probabilistic modeling method detailed in (8). For all analyses, cell matching occurred across two sessions and was performed using the following steps. First, centroid location for all cells were projected onto a single image. Slight rotation and translation difference between sessions were adjusted to achieve maximal cross-correlation between sessions. Probabilistic modeling was then employed to determine which model (centroid distance vs spatial correlation) was optimal for each set of data. For all data, the spatial correlation model yielded the best results and was thus used to match cells across sessions. For final alignment, the spatial correlation (not the joint model) was used, and correlation values for nearest neighbors was set individually for each animal depending on the intersection of the two models.

#### Encoding model

As described previously (9), a multiple linear regression was used to determine how behaviour contributed to neural activity. The dependent variable in the model was the z-scored, denoised calcium traces (F) of putative neurons identified by CMNFe.

For analysis of modulation in response to grooming, grooming was the only independent variable in the encoding model and was represented as a binary trace with a value of 1 when the animal was grooming and 0 elsewhere. The encoding model for grooming was:

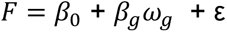

For analysis of modulation in response to reversal learning, each of the 5 behavioral events (correct lever presses (c), incorrect lever presses (i), magazine entries (e), reward cues (r), trial start cues (s)) were predictors in the model. Each predictor was represented as a vector of ‘1’s when events occurred and 0 elsewhere. Prior to being included in the model, each predictor was convolved with a 3s Gaussian.

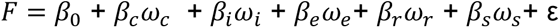

For both encoding models, F is the calcium trace for an individual neuron, ω is the behavior predictor, β is the regression coefficient, and ε is a Gaussian noise term. The β values were calculated using the least squares criterion.

To determine whether individual neurons were significantly modulated in response to each behavior variable, we ran two models: a full model containing all behavioral variables, and a partial model excluding the variable of interest. We next calculated an F-statistic for a nested model comparison between the partial model and the full model. We then created a null distribution of the same F-statistic. To do this we created 500 instances of a shuffled calcium trace by shuffling non-overlapping 3s bins to preserve the autocorrelation of the calcium trace. If the true F-statistic was greater than 3 standard deviations than the mean of the null distribution, then the neuron was considered significantly modulated in response to the behavior.

If a neuron was significantly modulated in response to the behavior, the strength of modulation was quantified using the regression coefficient (β).

#### Statistical analysis

Effects of genotype and fluoxetine on behavior were assessed using repeated measures ANOVA, except for bout length analysis which used Kolmogorov-Smirnov test to compare cumulative distribution frequencies between groups. For both grooming and reversal learning imaging analyses, effects of genotype and fluoxetine on percentages of cells modulated and strength of modulation were assessed using repeated measures ANOVA and linear regression, respectively.

Percentages of cells modulated in response to reversal learning were adjusted for differences in the number of behavioral events. Beta weights of cells modulated in response to reversal learning were adjusted for animal-to-animal variability and number of behavioral events. To make these adjustments, we performed a linear regression which included animal-to-animal variability and/or number of behavioral events as covariates. The mean value of the percentage of neurons (or beta-weight) across all groups was then added to the residuals of this regression. The result was then used as the dependent variable for the repeated measures ANOVA which analyzed the effects of genotype and drug. Percentage of cells and beta weights for grooming analyses were not adjusted for the number of grooming events because all KOs had a higher number of grooming bouts than WTs. Consequently, it was impossible to dissociate genotype effects from the effects of the number of grooming bouts.

#### Analysis of expected versus actual overlap between grooming and reversal learning

Data were pooled across all animals within genotype. To determine whether the actual overlap between reversal learning and grooming-modulated cells was different from what would be expected by chance, we performed a bootstrap analysis. For each of the 5 behaviors examined for reversal learning, we found the total number of neurons that are modulated in response to that behavior. As an example, for each genotype we calculated the total number of neurons modulated in response to the correct lever press (N_correct_) and the total number of neurons inhibited by grooming (N_groom_inh_). The actual overlap in the population is the percentage of neurons that are both modulated in response to the correct lever press and are inhibited by grooming. To construct the expected overlap, we randomly chose a sample of neurons (N_random_) equal to N_correct_ and calculated the percent overlap between N_random_ and N_groom_inh_. We repeated this procedure 1000 times to construct a distribution of the expected overlap between grooming and reversal learning populations. The 95% confidence interval of this distribution was represented as the shaded rectangles in Fig. 6,S10. We consider the populations to be overlapping, segregated or, randomly overlapping vs. segregated if the actual overlap is greater, less than or within the 95% CI of the expected overlap, respectively.

## Supplemental Discussion

### Type of neurons imaged

A variety of studies across species have shown that GABAergic interneurons make up between 10-30% of cortical neurons. We therefore predicted that 10-30% of cells in our dataset are GABAergic, under the assumption that the hSyn promotor expresses GCaMP equally well in excitatory and inhibitory neurons. To test whether this prediction was borne out in our imaging data, we determined whether we could detect distinct population clusters using two metrics shown to differ (10–12) between excitatory and inhibitory interneurons: 1) calcium event rate and 2) calcium event width. We chose event rate because cortical somatostatin (SOM+) interneurons, which comprise ∼30% of all GABAergic interneurons, have 2-3 times higher GCaMP6m event rates compared with cortical excitatory neurons (10). We chose event width because several prior studies have shown that GCaMP6f events in PV+ interneurons (11, 12), which make up ∼40% of cortical GABAergic neurons, are 3-4 times wider than events in excitatory pyramidal neurons.

We first analyzed imaging data from the pre-fluoxetine grooming (Fig.S2a-i) and reversal learning (Fig.S2j-r) sessions and plotted the distribution of event rates and event widths. Event times used to calculate event rate and width were identified by CNMFe. Event width was calculated by first averaging across all events from each neuron and then calculating the full width at half-max of the mean event trace.

For neurons recorded during the pre-fluoxetine grooming session, the distribution of event rates from all neurons did not differ between KO and WT neurons, and approximately 10% of cells from WTs and KOs had event rates greater than 2 times the median of 0.1Hz (Fig. S2a,d). Importantly, distributions of event rates from groom_exc_ (Fig. S2b,e) and groom_inh_ (Fig. S2c,f) neurons did not differ between WT and KOs. Approximately 5-10% of neurons had widths that are 3-4 times greater than the median width of 700ms (Fig. S2g,h), and similarly, the distribution of event widths did not differ between KO and WT neurons. Together, event rate and event width define a cluster of ∼10% of neurons (Fig. S2i) that are at least 2-3 fold different than the median of the entire population, suggesting that they are a distinct subset of neurons with characteristics that resemble GABAergic interneurons.

Similarly, for neurons recorded during the pre-fluoxetine reversal learning session, the distribution of event rates for all neurons did not differ between KO and WT neurons, and ∼10% of cells from both WTs and KOs had rates greater than twice the median of 0.1Hz (Fig. S2j,m). Additionally, distributions of event rates from neurons modulated in response to correct lever presses (Fig. S2k,n) or reward cues (Fig. S2l,o) did not differ between WT and KOs, and the distribution of event widths did not differ between KO and WT neurons (Fig. S2p,q). Finally, similar to the grooming session, event rate and width defined a cluster of ∼10% of neurons (Fig. S2r), suggesting that the size of the putative GABAergic neuron subset is similar during grooming and reversal learning sessions.

Because the proportion of putative GABAergic neurons is the same in KOs and WTs, we do not believe their presence significantly impacts our interpretation of results. For instance, one main conclusion is that decreased grooming in KOs after fluoxetine treatment is due (in part) to decreases in the percentage of groom_inh_ LOFC neurons. Because the same percentage of WT and KO groom_inh_ neurons belong to the putative GABAergic neuron cluster, the genotype difference in the percentage of groom_inh_ neurons is unlikely to be an artifact of imaging a different proportion of excitatory and inhibitory neurons across genotypes. Similarly, we found that decreased reversal learning performance in KOs is due, in part, to deficits in the strength of modulation of the correct lever press and reward cue. Because the same percentage of WT and KO neurons belong to the putative GABAergic neuron cluster, this genotype difference is unlikely to be an artifact of imaging a different proportion of excitatory and inhibitory neurons across genotypes.

### Effects of fluoxetine on grooming behavior

One reason why the Sapap3 KO mouse has become one of the most widely used rodent models for the study of compulsive behaviors is that, similar to patients with OCD, compulsive grooming improves after administration of an SSRI (13). Here we find that the number of grooming bouts decreases and the length of grooming bouts increases after fluoxetine suggesting that, at baseline, KOs have a disruption in the completion of grooming sequences. We find that after fluoxetine, both WTs and KOs display fewer bouts lasting less than 5 seconds and more bouts lasting greater than 5 seconds. Recent work showing that 3 seconds discriminates between short, single phase and longer, multi-phase grooming bouts adds evidence to the conclusion that fluoxetine allows mice to complete multi-phase grooming sequences (14).

### Role of the LOFC in fluoxetine-associated decreases in grooming

Here we provide evidence that fluoxetine decreases the number of grooming bouts in KOs by reducing the number of grooming inhibited LOFC neurons. While this evidence is indirect, our conclusions are supported by several lines of evidence. First, serotonin reuptake inhibitors (SRIs), the first line pharmacotherapy for patients with OCD, normalize orbitofrontal cortex activity in patients with OCD who respond clinically to treatment (15, 16). Second, optogenetic activation of the LOFC is sufficient to decrease grooming in KOs (17). Therefore, while fluoxetine is likely to exert numerous effects in multiple brain areas, decreases in grooming in KOs are likely caused, in part, by decreases in the number of grooming inhibited LOFC neurons.

### Role of the LOFC in fluoxetine-associated improvements in reversal learning

The role of the LOFC in reversal learning is complex. It is not required for reversal learning performance, as lesions to the LOFC that spare passing white matter tracts do not impair performance (18); however, the LOFC does play a role in updating the value of expected outcomes (19–21) and storing response-outcome associations (22). Additionally, the LOFC is likely to partially mediate fluoxetine-mediated improvements in reversal learning. Serotonin depletion in the OFC of rodents disrupts reversal learning (23), while either pharmacologic (24–26) or genetic (24) serotonin enhancement improves reversal learning. The data presented here, which show that fluoxetine-associated improvements in behavior parallel fluoxetine-associated improvements in modulation of LOFC neurons by task events, corroborate prior data and suggest that the behavior improvements with fluoxetine are mediated by changes in LOFC activity.

## References

1. Endrass T, Koehne S, Riesel A, Kathmann N (2013): Neural correlates of feedback processing in obsessive–compulsive disorder. Journal of Abnormal Psychology 122. https://doi.org/10.1037/a0031496

2. Remijnse PL, Nielen MMA, van Balkom AJLM, Cath DC, van Oppen P, Uylings HBM, Veltman DJ (2006): Reduced Orbitofrontal-Striatal Activity on a Reversal Learning Task in Obsessive-Compulsive Disorder. Archives of General Psychiatry 63. https://doi.org/10.1001/archpsyc.63.11.1225

3. Remijnse PL, Nielen MMA, van Balkom AJLM, Hendriks G-J, Hoogendijk WJ, Uylings HBM, Veltman DJ (2009): Differential frontal–striatal and paralimbic activity during reversal learning in major depressive disorder and obsessive–compulsive disorder. Psychological Medicine 39. https://doi.org/10.1017/S0033291708005072

4. Valerius G, Lumpp A, Kuelz A-K, Freyer T, Voderholzer U (2008): Reversal Learning as a Neuropsychological Indicator for the Neuropathology of Obsessive Compulsive Disorder? A Behavioral Study. The Journal of Neuropsychiatry and Clinical Neurosciences 20. https://doi.org/10.1176/jnp.2008.20.2.210

5. Szabó C, Németh A, Kéri S (2013): Ethical sensitivity in obsessive-compulsive disorder and generalized anxiety disorder: The role of reversal learning. Journal of Behavior Therapy and Experimental Psychiatry 44. https://doi.org/10.1016/j.jbtep.2013.04.001

6. Morein-Zamir S, Papmeyer M, Pertusa A, Chamberlain SR, Fineberg NA, Sahakian BJ, et al. (2014): The profile of executive function in OCD hoarders and hoarding disorder. Psychiatry Research 215. https://doi.org/10.1016/j.psychres.2013.12.026

7. Kim HW, Kang JI, Namkoong K, Jhung K, Ha RY, Kim SJ (2015): Further evidence of a dissociation between decision-making under ambiguity and decision-making under risk in obsessive–compulsive disorder. Journal of Affective Disorders 176. https://doi.org/10.1016/j.jad.2015.01.060

8. Chamberlain SR, Fineberg NA, Blackwell AD, Clark L, Robbins TW, Sahakian BJ (2007): A neuropsychological comparison of obsessive–compulsive disorder and trichotillomania. Neuropsychologia 45. https://doi.org/10.1016/j.neuropsychologia.2006.07.016

9. Remijnse PL, van den Heuvel OA, Nielen MMA, Vriend C, Hendriks G-J, Hoogendijk WJG, et al. (2013): Cognitive Inflexibility in Obsessive-Compulsive Disorder and Major Depression Is Associated with Distinct Neural Correlates. PLoS ONE 8. https://doi.org/10.1371/journal.pone.0059600

10. Chamberlain SR, Menzies L, Hampshire A, Suckling J, Fineberg NA, del Campo N, et al. (2008): Orbitofrontal Dysfunction in Patients with Obsessive-Compulsive Disorder and Their Unaffected Relatives. Science 321. https://doi.org/10.1126/science.1154433

11. Baxter LR (1987): Local Cerebral Glucose Metabolic Rates in Obsessive-Compulsive Disorder. Archives of General Psychiatry 44. https://doi.org/10.1001/archpsyc.1987.01800150017003

12. Rauch SL (1994): Regional Cerebral Blood Flow Measured During Symptom Provocation in Obsessive-Compulsive Disorder Using Oxygen 15—Labeled Carbon Dioxide and Positron Emission Tomography. Archives of General Psychiatry 51. https://doi.org/10.1001/archpsyc.1994.03950010062008

13. Breiter HC (1996): Functional Magnetic Resonance Imaging of Symptom Provocation in Obsessive-compulsive Disorder. Archives of General Psychiatry 53. https://doi.org/10.1001/archpsyc.1996.01830070041008

14. Welch JM, Lu J, Rodriguiz RM, Trotta NC, Peca J, Ding J-D, et al. (2007): Cortico-striatal synaptic defects and OCD-like behaviours in Sapap3-mutant mice. Nature 448. https://doi.org/10.1038/nature06104

15. Burguière E, Monteiro P, Feng G, Graybiel AM (2013): Optogenetic Stimulation of Lateral Orbitofronto-Striatal Pathway Suppresses Compulsive Behaviors. Science 340. https://doi.org/10.1126/science.1232380

16. Ade KK, Wan Y, Hamann HC, O’Hare JK, Guo W, Quian A, et al. (2016): Increased Metabotropic Glutamate Receptor 5 Signaling Underlies Obsessive-Compulsive Disorder-like Behavioral and Striatal Circuit Abnormalities in Mice. Biological Psychiatry 80. https://doi.org/10.1016/j.biopsych.2016.04.023

17. Wan Y, Ade KK, Caffall Z, Ilcim Ozlu M, Eroglu C, Feng G, Calakos N (2014): Circuit-Selective Striatal Synaptic Dysfunction in the Sapap3 Knockout Mouse Model of Obsessive-Compulsive Disorder. Biological Psychiatry 75. https://doi.org/10.1016/j.biopsych.2013.01.008

18. Welch JM, Wang D, Feng G (2004): Differential mRNA expression and protein localization of the SAP90/PSD-95-associated proteins (SAPAPs) in the nervous system of the mouse. The Journal of Comparative Neurology 472. https://doi.org/10.1002/cne.20060

19. Züchner S, Wendland JR, Ashley-Koch AE, Collins AL, Tran-Viet KN, Quinn K, et al. (2009): Multiple rare SAPAP3 missense variants in trichotillomania and OCD. Molecular Psychiatry 14. https://doi.org/10.1038/mp.2008.83

20. Mattheisen M, Samuels JF, Wang Y, Greenberg BD, Fyer AJ, McCracken JT, et al. (2015): Genome-wide association study in obsessive-compulsive disorder: results from the OCGAS. Molecular Psychiatry 20. https://doi.org/10.1038/mp.2014.43

21. Stewart SE, Yu D, Scharf JM, Neale BM, Fagerness JA, Mathews CA, et al. (2013): Genome-wide association study of obsessive-compulsive disorder. Molecular Psychiatry 18. https://doi.org/10.1038/mp.2012.85

22. Bienvenu OJ, Wang Y, Shugart YY, Welch JM, Grados MA, Fyer AJ, et al. (2009): *Sapap3* and pathological grooming in humans: Results from the OCD collaborative genetics study. American Journal of Medical Genetics Part B: Neuropsychiatric Genetics 150B. https://doi.org/10.1002/ajmg.b.30897

23. Piantadosi SC, Chamberlain BL, Glausier JR, Lewis DA, Ahmari SE (2021): Lower excitatory synaptic gene expression in orbitofrontal cortex and striatum in an initial study of subjects with obsessive compulsive disorder. Molecular Psychiatry 26. https://doi.org/10.1038/s41380-019-0431-3

24. Manning EE, Dombrovski AY, Torregrossa MM, Ahmari SE (2019): Impaired instrumental reversal learning is associated with increased medial prefrontal cortex activity in Sapap3 knockout mouse model of compulsive behavior. Neuropsychopharmacology 44. https://doi.org/10.1038/s41386-018-0307-2

25. Benzina N, N’Diaye K, Pelissolo A, Mallet L, Burguière E (2021): A cross-species assessment of behavioral flexibility in compulsive disorders. Communications Biology 4. https://doi.org/10.1038/s42003-020-01611-y

26. Yang Z, Wu G, Liu M, Sun X, Xu Q, Zhang C, Lei H (2021): Dysfunction of Orbitofrontal GABAergic Interneurons Leads to Impaired Reversal Learning in a Mouse Model of Obsessive-Compulsive Disorder. Current Biology 31. https://doi.org/10.1016/j.cub.2020.10.045

27. van den Boom BJG, Mooij AH, Misevičiūtė I, Denys D, Willuhn I (2019): Behavioral flexibility in a mouse model for obsessive-compulsive disorder: Impaired Pavlovian reversal learning in SAPAP3 mutants. Genes, Brain and Behavior 18. https://doi.org/10.1111/gbb.12557

28. Resendez SL, Jennings JH, Ung RL, Namboodiri VMK, Zhou ZC, Otis JM, et al. (2016): Visualization of cortical, subcortical and deep brain neural circuit dynamics during naturalistic mammalian behavior with head-mounted microscopes and chronically implanted lenses. Nature Protocols 11. https://doi.org/10.1038/nprot.2016.021

29. Corbit VL, Piantadosi SC, Wood J, Liu G, Choi CJY, Witten IB, et al. (2020): Dissociable roles of central striatum and anterior lateral motor area in initiating and sustaining naturalistic behavior. bioRxiv 2020.01.08.899070.

30. Ahmari SE, Spellman T, Douglass NL, Kheirbek MA, Simpson HB, Deisseroth K, et al. (2013): Repeated Cortico-Striatal Stimulation Generates Persistent OCD-Like Behavior. Science 340. https://doi.org/10.1126/science.1234733

31. Dulawa SC, Holick KA, Gundersen B, Hen R (2004): Effects of Chronic Fluoxetine in Animal Models of Anxiety and Depression. Neuropsychopharmacology 29. https://doi.org/10.1038/sj.npp.1300433

32. Zhou P, Resendez SL, Rodriguez-Romaguera J, Jimenez JC, Neufeld SQ, Giovannucci A, et al. (2018): Efficient and accurate extraction of in vivo calcium signals from microendoscopic video data. eLife 7. https://doi.org/10.7554/eLife.28728

33. Sheintuch L, Rubin A, Brande-Eilat N, Geva N, Sadeh N, Pinchasof O, Ziv Y (2017): Tracking the Same Neurons across Multiple Days in Ca2+ Imaging Data. Cell Reports 21. https://doi.org/10.1016/j.celrep.2017.10.013

34. Engelhard B, Finkelstein J, Cox J, Fleming W, Jang HJ, Ornelas S, et al. (2019): Specialized coding of sensory, motor and cognitive variables in VTA dopamine neurons. Nature 570. https://doi.org/10.1038/s41586-019-1261-9

35. Stalnaker TA, Cooch NK, Schoenbaum G (2015): What the orbitofrontal cortex does not do. Nature Neuroscience 18. https://doi.org/10.1038/nn.3982

36. Izquierdo A, Brigman JL, Radke AK, Rudebeck PH, Holmes A (2017): The neural basis of reversal learning: An updated perspective. Neuroscience 345. https://doi.org/10.1016/j.neuroscience.2016.03.021

37. Alsiö J, Lehmann O, McKenzie C, Theobald DE, Searle L, Xia J, et al. (2021): Serotonergic Innervations of the Orbitofrontal and Medial-prefrontal Cortices are Differentially Involved in Visual Discrimination and Reversal Learning in Rats. Cerebral Cortex 31. https://doi.org/10.1093/cercor/bhaa277

38. Robbins TW, Vaghi MM, Banca P (2019): Obsessive-Compulsive Disorder: Puzzles and Prospects. Neuron 102. https://doi.org/10.1016/j.neuron.2019.01.046

## References

1. Manning EE, Dombrovski AY, Torregrossa MM, Ahmari SE (2019): Impaired instrumental reversal learning is associated with increased medial prefrontal cortex activity in Sapap3 knockout mouse model of compulsive behavior. Neuropsychopharmacology 44. https://doi.org/10.1038/s41386-018-0307-2

2. Ahmari SE, Spellman T, Douglass NL, Kheirbek MA, Simpson HB, Deisseroth K, et al. (2013): Repeated Cortico-Striatal Stimulation Generates Persistent OCD-Like Behavior. Science 340. https://doi.org/10.1126/science.1234733

3. Dulawa SC, Holick KA, Gundersen B, Hen R (2004): Effects of Chronic Fluoxetine in Animal Models of Anxiety and Depression. Neuropsychopharmacology 29. https://doi.org/10.1038/sj.npp.1300433

4. Kalueff A v., Stewart AM, Song C, Berridge KC, Graybiel AM, Fentress JC (2016): Neurobiology of rodent self-grooming and its value for translational neuroscience. Nature Reviews Neuroscience 17. https://doi.org/10.1038/nrn.2015.8

5. Ghosh KK, Burns LD, Cocker ED, Nimmerjahn A, Ziv Y, Gamal A el, Schnitzer MJ (2011): Miniaturized integration of a fluorescence microscope. Nature Methods 8. https://doi.org/10.1038/nmeth.1694

6. Zhou P, Resendez SL, Rodriguez-Romaguera J, Jimenez JC, Neufeld SQ, Giovannucci A, et al. (2018): Efficient and accurate extraction of in vivo calcium signals from microendoscopic video data. eLife 7. https://doi.org/10.7554/eLife.28728

7. Resendez SL, Jennings JH, Ung RL, Namboodiri VMK, Zhou ZC, Otis JM, et al. (2016): Visualization of cortical, subcortical and deep brain neural circuit dynamics during naturalistic mammalian behavior with head-mounted microscopes and chronically implanted lenses. Nature Protocols 11. https://doi.org/10.1038/nprot.2016.021

8. Sheintuch L, Rubin A, Brande-Eilat N, Geva N, Sadeh N, Pinchasof O, Ziv Y (2017): Tracking the Same Neurons across Multiple Days in Ca2+ Imaging Data. Cell Reports 21. https://doi.org/10.1016/j.celrep.2017.10.013

9. Engelhard B, Finkelstein J, Cox J, Fleming W, Jang HJ, Ornelas S, et al. (2019): Specialized coding of sensory, motor and cognitive variables in VTA dopamine neurons. Nature 570. https://doi.org/10.1038/s41586-019-1261-9

10. Scheggia D, Managò F, Maltese F, Bruni S, Nigro M, Dautan D, et al. (2020): Somatostatin interneurons in the prefrontal cortex control affective state discrimination in mice. Nature Neuroscience 23. https://doi.org/10.1038/s41593-019-0551-8

11. Gritton HJ, Howe WM, Romano MF, DiFeliceantonio AG, Kramer MA, Saligrama V, et al. (2019): Unique contributions of parvalbumin and cholinergic interneurons in organizing striatal networks during movement. Nature Neuroscience 22. https://doi.org/10.1038/s41593-019-0341-3

12. Pinto L, Dan Y (2015): Cell-Type-Specific Activity in Prefrontal Cortex during Goal-Directed Behavior. Neuron 87. https://doi.org/10.1016/j.neuron.2015.06.021

13. Welch JM, Lu J, Rodriguiz RM, Trotta NC, Peca J, Ding J-D, et al. (2007): Cortico-striatal synaptic defects and OCD-like behaviours in Sapap3-mutant mice. Nature 448. https://doi.org/10.1038/nature06104

14. Lamothe H, Schreiweis C, Lavielle O, Mallet L, Burguière E (2021): Not only compulsivity: The SAPAP3-KO mouse reconsidered as a comorbid model expressing a spectrum of pathological repetitive behaviors. bioRxiv 2020.01.22.915215.

15. van der Straten AL, Denys D, van Wingen GA (2017): Impact of treatment on resting cerebral blood flow and metabolism in obsessive compulsive disorder: a meta-analysis. Scientific Reports 7. https://doi.org/10.1038/s41598-017-17593-7

16. Benkelfat C (1990): Local Cerebral Glucose Metabolic Rates in Obsessive-Compulsive Disorder. Archives of General Psychiatry 47. https://doi.org/10.1001/archpsyc.1990.01810210048007

17. Burguière E, Monteiro P, Feng G, Graybiel AM (2013): Optogenetic Stimulation of Lateral Orbitofronto-Striatal Pathway Suppresses Compulsive Behaviors. Science 340. https://doi.org/10.1126/science.1232380

18. Rudebeck PH, Saunders RC, Prescott AT, Chau LS, Murray EA (2013): Prefrontal mechanisms of behavioral flexibility, emotion regulation and value updating. Nature Neuroscience 16. https://doi.org/10.1038/nn.3440

19. Stalnaker TA, Cooch NK, Schoenbaum G (2015): What the orbitofrontal cortex does not do. Nature Neuroscience 18. https://doi.org/10.1038/nn.3982

20. Bissonette GB, Martins GJ, Franz TM, Harper ES, Schoenbaum G, Powell EM (2008): Double Dissociation of the Effects of Medial and Orbital Prefrontal Cortical Lesions on Attentional and Affective Shifts in Mice. Journal of Neuroscience 28. https://doi.org/10.1523/JNEUROSCI.2820-08.2008

21. Schoenbaum G, Chiba AA, Gallagher M (2000): Changes in Functional Connectivity in Orbitofrontal Cortex and Basolateral Amygdala during Learning and Reversal Training. The Journal of Neuroscience 20. https://doi.org/10.1523/JNEUROSCI.20-13-05179.2000

22. Keiflin R, Reese RM, Woods CA, Janak PH (2013): The Orbitofrontal Cortex as Part of a Hierarchical Neural System Mediating Choice between Two Good Options. Journal of Neuroscience 33. https://doi.org/10.1523/JNEUROSCI.0026-13.2013

23. Alsiö J, Lehmann O, McKenzie C, Theobald DE, Searle L, Xia J, et al. (2021): Serotonergic Innervations of the Orbitofrontal and Medial-prefrontal Cortices are Differentially Involved in Visual Discrimination and Reversal Learning in Rats. Cerebral Cortex 31. https://doi.org/10.1093/cercor/bhaa277

24. Brigman JL, Mathur P, Harvey-White J, Izquierdo A, Saksida LM, Bussey TJ, et al. (2010): Pharmacological or Genetic Inactivation of the Serotonin Transporter Improves Reversal Learning in Mice. Cerebral Cortex 20. https://doi.org/10.1093/cercor/bhp266

25. Bari A, Theobald DE, Caprioli D, Mar AC, Aidoo-Micah A, Dalley JW, Robbins TW (2010): Serotonin Modulates Sensitivity to Reward and Negative Feedback in a Probabilistic Reversal Learning Task in Rats. Neuropsychopharmacology 35. https://doi.org/10.1038/npp.2009.233

26. Zhukovsky P, Alsiö J, Jupp B, Xia J, Guiliano C, Jenner L, et al. (2017): Perseveration in a spatial-discrimination serial reversal learning task is differentially affected by MAO-A and MAO-B inhibition and associated with reduced anxiety and peripheral serotonin levels. Psychopharmacology 234. https://doi.org/10.1007/s00213-017-4569-x

