## Supplementary material for "Independent and distinct patterns of abnormal lateral orbitofrontal cortex activity during compulsive grooming and reversal learning normalize after fluoxetine": Table 3

**FIGURE 1**

| | Avg ± sem WT Pre | WT Post | KO Pre | KO Post | Initial Repeated measures ANOVA (p-values) Genotype | Drug | Interaction | Post-hoc t-test (p values from unpaired ttest unless indicated) WT Pre vs Post (paired ttest) | WT Pre vs KO Pre | WT Pre vs KO Post | WT Post vs KO Pre | WT Post vs KO Post | KO Pre vs KO Post (paired ttest) |
|---|---|---|---|---|---|---|---|---|---|---|---|---|---|
| 1c: Grooming Time | 9.8 ± 1.6 | 11.7 ± 1.7 | 42.1 ± 11.8 | 38.9 ± 12.5 | 0.01 | 0.08 | 0.88 | 0.19 | < 0.01 | < 0.01 | < 0.01 | < 0.01 | 0.2 |
| 1d: Grooming Bouts | 42.0 ± 5.0 | 40.8 ± 6.2 | 159.2 ± 25.6 | 107.0 ± 24.0 | 0.003 | 0.01 | 0.03 | 0.25 | < 0.01 | < 0.01 | < 0.01 | 0.013 | 0.02 |

| | KS test p-value |
|---|---|
| 1e: WT vs KO bout length CDF | < 10e-5 |
| 1f: WT pre vs post bout length CDF | < 10e-5 |
| 1g: KO pre vs post bout length CDF | <10e-5 |

**FIGURE 2**

| | Avg ± sem WT Pre | WT Post | KO Pre | KO Post | Initial Repeated measures ANOVA (p-values) Genotype | Drug | Interaction | Post-hoc t-test (p values from unpaired ttest unless indicated) WT Pre vs Post (paired ttest for 2f-g) | WT Pre vs KO Pre | WT Pre vs KO Post | WT Post vs KO Pre | WT Post vs KO Post | KO Pre vs KO Post (paired ttest for 2fg) |
|---|---|---|---|---|---|---|---|---|---|---|---|---|---|
| 2f: Percent inhibited cells | 22.1 ± 3.6 | 20.9 ± 4.9 | 32.1 ± 1.6 | 25.3 ± 1.8 | 0.02 | < 0.001 | < 0.001 | 0.68 | < 0.01 | 0.42 | < 0.01 | 0.45 | 0.01 |
| 2g: Percent excited cells | 26.8 ± 6.0 | 26.5 ± 8.6 | 27.1 ± 6.0 | 27.4 ± 6.7 | 82 | 0.78 | 0.96 | 0.98 | 0.76 | 0.72 | 0.88 | 0.9 | 0.87 |
| 2h: Beta inhibited cells | 0.08 ± 0.01 | 0.08 ± 0.02 | 0.16 ± 0.01 | 0.15 ± 0.01 | <0.001 | 0.67 | 0.59 | 0.94 | <0.01 | <0.01 | <0.01 | <0.01 | 0.89 |
| 2i: Beta excited cells | 0.14 ± 0.02 | 0.14 ± 0.02 | 0.18 ± 0.01 | 0.18 ± 0.01 | <0.001 | 0.61 | 0.52 | 0.99 | <0.01 | <0.01 | <0.01 | <0.01 | 0.94 |

| | Correlation r WT | KO | Correlation p-value WT | KO |
|---|---|---|---|---|
| 2j: bouts vs inhibited cells | 0.54 | 0.25 | 0.52 | 0.24 |
| 2k: change bouts vs change inhib | -0.4 | 0.82 | 0.41 | 0.01 |

**FIGURE 3**

| | Avg ± sem WT Pre | WT Post | KO Pre | KO Post | Initial Repeated measures ANOVA (p-values) Genotype | Drug | Interaction | Post-hoc t-test (p values from unpaired ttest unless indicated) WT Pre vs Post (paired ttest) | WT Pre vs KO Pre | WT Pre vs KO Post | WT Post vs KO Pre | WT Post vs KO Post | KO Pre vs KO Post (paired ttest) |
|---|---|---|---|---|---|---|---|---|---|---|---|---|---|
| 3b: Percent inhibited cells | 22.5 ± 4.1 | 23.6 ± 2.4 | 29.9 ± 2.3 | 24.8 ± 1.5 | 0.04 | 0.03 | 0.04 | 0.67 | 0.03 | 0.23 | 0.02 | 0.19 | 0.04 |

| | Avg ± sem WT | KO | t-test pvalue |
|---|---|---|---|
| 3c: change in inhibited cells | 110 ± 11 | 83 ± 4 | 0.02 |
| 3d: remaining inhibited cells | 41 ± 12 | 45 ± 6 | 0.54 |
| 3e: newly inhibited cells | 67 ± 3 | 38 ± 5 | <0.001 |

**FIGURE 4**

| | Avg ± sem WT Pre | WT Post | KO Pre | KO Post | Initial Repeated measures ANOVA (p-values) Genotype | Drug | Interaction | Post-hoc t-test (p values from unpaired ttest unless indicated) WT Pre vs Post (paired ttest) | WT Pre vs KO Pre | WT Pre vs KO Post | WT Post vs KO Pre | WT Post vs KO Post | KO Pre vs KO Post (paired ttest) |
|---|---|---|---|---|---|---|---|---|---|---|---|---|---|
| 4c: Correct Presses | 68 ± 7 | 59 ± 7 | 33 ± 10 | 59 ± 8 | 0.03 | 0.39 | 0.04 | 0.16 | 0.02 | 0.34 | 0.04 | 0.85 | 0.02 |
| 4e: Correct mid point | 7.4 ± 0.8 | 7.1 ± 0.4 | 11.1 ± 1.2 | 7. 4 ± 0.4 | 0.008 | 0.25 | 0.02 | 0.45 | 0.02 | 0.65 | 0.01 | 0.81 | 0.01 |
| 4f: Reward cue | 38.1 ± 3.9 | 29.2 ± 4.0 | 15.9 ± 5.0 | 29.5 ± 4.1 | 0.02 | 0.82 | 0.02 | 0.03 | < 0.01 | 0.1 | 0.05 | 0.51 | 0.04 |
| 4g: Incorrect presses | 72 ± 10 | 35 ± 5 | 88 ± 20 | 74 ± 18 | 0.3 | 0.01 | 0.15 | 0.01 | 0.19 | 0.65 | 0.01 | 0.04 | 0.04 |

**FIGURE 5**

| | Avg ± sem WT Pre | WT Post | KO Pre | KO Post | Initial Repeated measures ANOVA (p-values) Genotype | Drug | Interaction | Post-hoc t-test (p values from unpaired ttest unless indicated) WT Pre vs Post (paired ttest for 5e-g) | WT Pre vs KO Pre | WT Pre vs KO Post | WT Post vs KO Pre | WT Post vs KO Post | KO Pre vs KO Post (paired ttest for 5e-g) |
|---|---|---|---|---|---|---|---|---|---|---|---|---|---|
| 5e: Adj percent correct presses | 11.8 ± 1.2 | 10.2 ± 2.2 | 9.0 ± 1.3 | 9.9 ± 2.6 | 0.38 | 0.78 | 0.27 | 0.77 | 0.06 | 0.42 | 0.19 | 0.89 | 0.28 |
| 5f: Adj percent reward cue | 6.8 ± 0.6 | 7.7 ± 0.6 | 6.0 ± 0.4 | 4.1 ± 0.7 | 0.24 | 0.6 | 0.26 | 0.34 | 0.55 | 0.13 | 0.48 | 0.09 | 0.13 |
| 5g: Adj percent incorrect presses | 19.8 ± 1.4 | 14.5 ± 2.0 | 14.1 ± 2.2 | 14.2 ± 2.2 | 0.16 | 0.18 | 0.14 | 0.02 | 0.03 | 0.03 | 0.71 | 0.74 | 0.8 |
| 5h: Adj beta correct presses | 0.33 ± 0.03 | 0.30 ± 0.03 | 0.27 ± 0.02 | 0.30 ± 0.01 | 0.01 | 0.18 | 0.04 | 0.28 | 0.01 | 0.16 | 0.04 | 0.55 | 0.04 |
| 5i: Adj beta reward cue | 0.17 ± 0.02 | 0.16 ± 0.01 | 0.10 ± 0.01 | 0.14 ± 0.01 | 0.001 | 0.45 | 0.01 | 0.56 | <0.01 | 0.18 | < 0.01 | 0.39 | 0.01 |
| 5j: Adj beta incorrect presses | 0.33 ± 0.01 | 0.28 ± 0.01 | 0.33 ± 0.02 | 0.26 ± 0.01 | 0.65 | 0.01 | 0.43 | 0.02 | 0.71 | 0.02 | 0.04 | 0.19 | 0.02 |

**FIGURE 7**

| | Avg ± sem WT Pre | WT Post | KO Pre | KO Post | Initial Repeated measures ANOVA (p-values) Genotype | Drug | Interaction | Post-hoc t-test (p values from unpaired ttest unless indicated) WT Pre vs Post | WT Pre vs KO Pre | WT Pre vs KO Post | WT Post vs KO Pre | WT Post vs KO Post | KO Pre vs KO Post |
|---|---|---|---|---|---|---|---|---|---|---|---|---|---|
| 7d: Adj beta correct lever press | 0.22 ± 0.01 | 0.22 ± 0.01 | 0.13 ± 0.01 | 0.20 ± 0.01 | < 0.001 | 0.18 | < 0.001 | 0.89 | < 0.01 | 0.71 | < 0.01 | 0.74 | 0.01 |
| 7g: Adj beta reward cue | 0.22 ± 0.02 | 0.20 ± 0.01 | 0.17 ± 0.01 | 0.20 ± 0.01 | < 0.01 | 0.45 | < 0.001 | 0.19 | 0.01 | 0.32 | 0.02 | 0.48 | 0.02 |

| | Avg ± sem WT No groom mod | WT $Groom_{exc}$ | WT $Groom_{inh}$ | KO No groom mod | KO $Groom_{exc}$ | KO $Groom_{inh}$ | Initial Repeated measures ANOVA (p-values) Genotype | Groom modulation | Interaction | Post-hoc t-test (p values from unpaired ttest unless indicated) No mod: WT vs KO | $Groom_{exc}$ : WT vs KC | $Groom_{inh}$ : WT vs K | WT: No mod vs Inh | KO: No mod vs Inh |
|---|---|---|---|---|---|---|---|---|---|---|---|---|---|---|
| 7e: Adj beta correct lever press pre | 0.23 ± 0.01 | 0.2 ± 0.02 | 0.19 ± 0.03 | 0.18 ± 0.01 | 0.16 ± 0.01 | 0.10 ± 0.01 | < 0.001 | 0.01 | 0.56 | < 0.01 | 0.01 | 0.02 | 0.05 | 0.01 |
| 7f: Adj beta correct lever press pos | 0.23 ± 0.02 | 0.19 ± 0.01 | 0.02 ± 0.01 | 0.023 ± 0.01 | 0.21 ± 0.01 | 0.18 ± 0.01 | 0.67 | 0.01 | 0.34 | 0.67 | 0.12 | 0.09 | 0.03 | <0.01 |
| 7h: Adj beta reward cue pre | 0.22 ± 0.02 | 0.23 ± 0.03 | 0.18 ± 0.01 | 0.19 ± 0.01 | 0.17 ± 0.01 | 0.16 ± 0.01 | 0.01 | 0.04 | 0.85 | 0.05 | 0.03 | 0.04 | 0.04 | < 0.01 |
| 7i: Adj beta reward cue post | 0.20 ± 0.01 | NA | 0.19 ± 0.010 | 0.22 ± 0.01 | 0.19 ± 0.01 | 0.19 ± 0.01 | 0.81 | 0.03 | 0.73 | 0.11 | NA | 0.54 | 0.04 | 0.01 |
